# Discovery of anthelmintic small molecules in the Medicines for Malaria Venture’s COVID and Global Health Priority Boxes using an infrared-based assay for *Caenorhabditis elegans* motility

**DOI:** 10.1101/2023.12.09.570935

**Authors:** Yujie Uli Sun, Lawrence J. Liu, Conor R. Caffrey

## Abstract

**Background:** Parasitic nematodes are a public health problem globally, and an economic burden on animal and plant agricultural industries. With their ability to generate drug resistance, new anthelmintic compounds must be constantly sourced.

**Methods:** Using the free-living nematode, *Caenorhabditis elegans,* in an infrared-based motility assay, we screened 400 compounds from two open-source, small-molecule collections distributed by the Medicines for Malaria Venture, namely, the COVID Box and Global Health Priority Box. The screening assay was first validated for worm number, DMSO concentration and final volume.

**Results:** Primary and secondary (time- and concentration-dependent) screens of both boxes, identified twelve compounds as hits; nine of which were known anthelmintics. Three novel anthelmintic hits, flufenerim, flucofuron and indomethacin were identified with EC_50_ values ranging from 0.211 to 23.174 µM. Counter toxicity screens with HEK293 cells indicated varying degrees of toxicity with EC_50_ values ranging from 0.453 to >100 µM.

**Conclusions:** A *C. elegans* motility assay was optimized and used to screen two recently-released, small molecule libraries. One or more of these three novel active compounds might serve as starting points for the development of new anthelmintics.

## Introduction

Parasitic nematodes infect more than one quarter of the global population [**Error! Bookmark not defined.**, **Error! Reference source not found.**] and promote poverty by constraining personal and societal productivity [2, 3]. Just a handful of drug classes are available [4], and failed treatments are known [5, 6, 7]. Nematode parasites are also an economic burden to agriculture [8–10] with drug resistance being quick to emerge [11]. Accordingly, new nematicides are urgently needed.

The free-living nematode, *Caenorhabditis elegans,* is useful in nematicide drug discovery [12–15]. We developed and deployed a *C. elegans* screen to evaluate two small molecule ‘box’ collections released by the drug development consortium, the Medicines for Malaria Venture (MMV). These COVID and Global Health Priority (GHP) Boxes, comprise 160 and 240 compounds, respectively, and contain bioactives for the SARS-CoV-2 virus, and various pathogens and vectors, respectively. Both boxes are the latest in a series of boxes distributed *gratis* by the MMV to spur drug discovery for infectious diseases [16–24].

## Methods

The COVID and GHP Boxes were provided by the MMV, Geneva, Switzerland as 10 µL solutions in DMSO (10 or 2 mM) in 96-well polypropylene plates. Plates were stored at – 80 °C.

Ivermectin, doramectin, selamectin and tolfenpyrad were from Sigma Aldrich. Milbemectin was from Cayman Chemical, and moxidectin, abamectin, chlorfenapyr, eprinomectin and indomethacin were from AK Scientific. Flucofuron was from ChemScene.

### *C. elegans* maintenance and motility assay

Methods to cultivate *C. elegans* (Bristol N2) and synchronize growth to the L4 stage were as described [25]. WMicroTracker ONE (Phylumtech, Argentina) detects movement by measuring the scattering of infrared light beams that are projected into each well of a microtiter plate (two 880 nm beams/well in a 96w plate) [26].

Synchronized *C. elegans* L4 were detached from agar and collected in M9 buffer. Worms were centrifuged for 1 min at 1,900 *g* and washed in S medium to decrease the concentration of *E. coli* that might interfere with infrared detection. Control and assay compounds (1 µL) in DMSO were spotted into each well of a clear, flat-bottomed 96-well polystyrene plate (Fisherbrand). DMSO was the negative control. Approximately 70 L4 in 100 µL S medium were added per well. First pass screens employed 40 µM compound, and motility was measured every 20 min for 24 h. Motility was normalized relative to the DMSO controls. Compounds were categorized according to the percentage motility remaining: 0 – 25% = potent; 26 – 65% = moderate; and >65% = low.

For potent compounds, concentration response assays (nine concentrations: 0.005 µM to 40 or 100 µM) were conducted to measure the half-maximal effective concentration (EC_50_). Compounds were serially diluted in DMSO using 96-well, v-bottomed polypropylene dilution plates, and 1 µL aliquots spotted into each well of the assay plates. Motility was measured as described above. Prism GraphPad, Version 8.0 (GraphPad Software, San Diego, CA) was used to calculate EC_50_ values using a non-linear sigmoidal four parameter logistic curve.

### Human embryonic kidney (HEK) 293 cytotoxicity assay

Potent compounds were also evaluated for cytotoxicity against HEK293 cells. Cells were cultured at 37 °C and 5% in DMEM containing 10% heat-inactivated FBS and 1% penicillin-streptomycin, and then sub-cultured when 60-80% confluent. Cells were detached with 0.05% trypsin/EDTA (Thermo Fisher Scientific) and centrifuged for 5 min at 1,900 *g*.

Screens were performed over 11 compound concentrations (0.00007 – 40 µM) that had been prepared in DMSO. Aliquots (1 µL) were spotted into each well of a clear-bottomed, 96-well, polystyrene assay plate (Fisherbrand). Approximately 20,000 HEK293 cells in 99 µL of the above supplemented DMEM were then added to each well. After 46 h at 37 °C and 5% CO_2_, 20 µL 0.5 mM resazurin (Alfa Aesar) were added and the incubation continued for 2 h at 37 °C [27]. Fluorescence was measured in a 2104 EnVision Multilabel Plate Reader (PerkinElmer) at 560 and 590 nm excitation and emission wavelengths, respectively. Raw data were exported from the EnVision Workstation software (version 1.13.3009.1401; PerkinElmer) into Prism GraphPad, and the half-maximal cytotoxic concentration (CC_50_) values were calculated using a non-linear regression curve.

## Results

### WMicroTracker assay optimization

To optimize the WMicroTracker assay, these variables were considered: the number of L4, the presence and concentration of DMSO, and the final volume per well.

Regarding the number of L4 per well, 30, 50, 60, 70, 80, 100, 150 and 200 L4 were tested in 100 µL S medium in the presence or absence of 1% DMSO. No significant effect was measured with or without DMSO, and for both conditions, motility trended upwards with increasing worm numbers (**Fig. 1A**). The greatest number of worms (150 or 200) resulted in the highest raw motility units, which would potentially improve the dynamic range of the assay; however, using so many worms would constrain assay throughput. By contrast, there was no statistically significant difference in the motility measured between 70 and 100 worms, and, in the interests of economy, we selected 70 L4 per well.

**Fig. 1.**
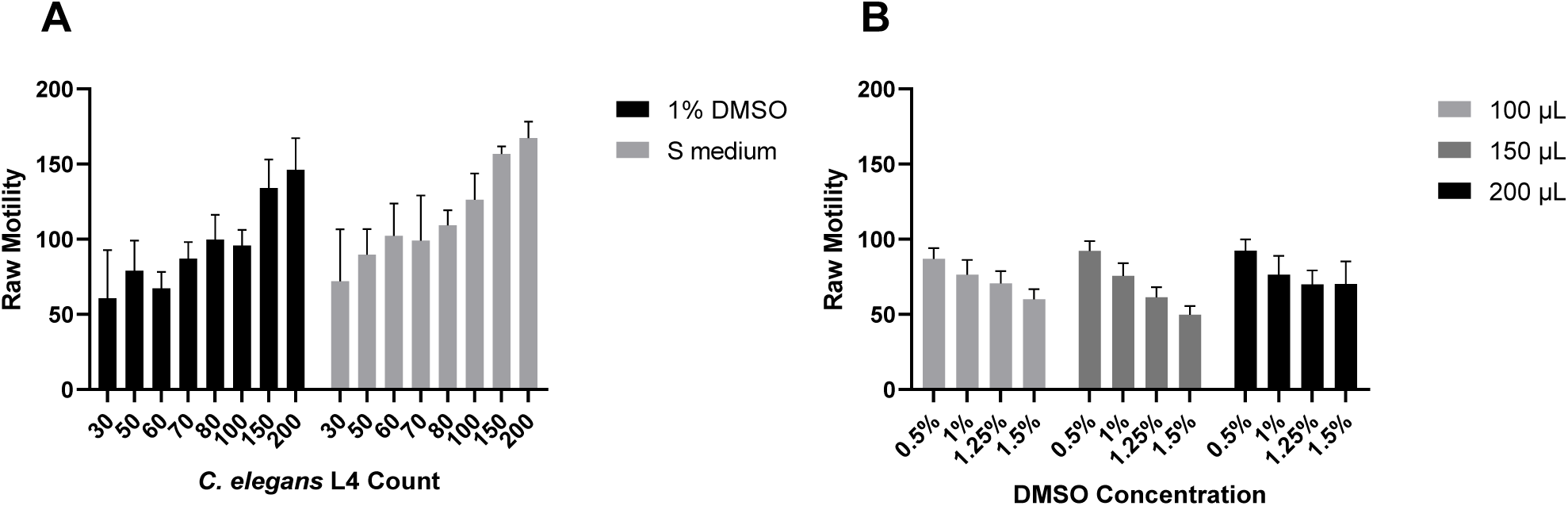
WMicroTracker assay optimization. (**A**) *C. elegans* motility as a function of worm number and the presence or absence of 1% DMSO in a final volume of 100 µL. The columns and error bars display the mean ± SD values from one experiment performed in triplicate. A one-sample *t*-test found no significant difference in worm motility in the presence or absence of DMSO at each of the final worm counts tested. A one-way ANOVA test found no significant difference in motility between 70 and 100 worms per well in either the presence or absence of DMSO. (**B**) *C. elegans* motility as a function of DMSO concentration and final assay volume. The columns and error bars display the mean ± SD values from one experiment performed in triplicate. A one-sample *t*-test found no difference between 0.5% and 1% DMSO for each of the three volumes tested. For each of the four DMSO concentrations tested, a one-sample *t*-test found no difference between the three volumes tested. For both **A** and **B**, motility in raw units was recorded at the 24 h time point.

Due to its thick collagen-enriched cuticle and extensive xenobiotic metabolism [28, 29], *C. elegans* is often exposed to relatively high concentrations of screen compounds. This requires consideration of the final concentration of DMSO to maintain solubility. Therefore, the effect of DMSO concentrations between 0.5 and 1.5 % on motility was measured. In parallel, we also considered the final assay volume (100, 150 and 200 µL).

Increasing the DMSO concentration decreased L4 motility in the final volumes of 100 and 150 µL, but less so in 200 µL (**Fig. 1B**). In all three volumes, motility at 0.5% and 1% DMSO showed no significant difference. Thus, a final concentration of 1% DMSO in 100 µL was chosen to maximize compound solubility in the least assay volume.

### Primary screening of the COVID and Global Health Priority Boxes

The COVID and GHP Box compounds were screened at 40 µM over 24 h (**Additional Table 1**). Twelve compounds were potent (motility <25% of DMSO) (**Fig. 2A**) with eleven of these from the GHP Box. Seven compounds are established macrocyclic lactone anthelmintics [30]: milbemectin (MMV1578924; 1.69% motility relative to control), moxidectin (MMV1633828; 0.28%), doramectin (MMV1633823; 6.74%), ivermectin (MMV672931; 3.37%), abamectin (MMV1577454; 3.60%), selamectin (MMV002231; 2.31%) and eprinomectin (MMV1633829; 13.19%). Another hit, the insecticide, tolfenpyrad (MMV688934; 0.26%), is a known active against the parasitic nematode, *Haemonchus contortus* [24] and *C. elegans* in screens of the MMV’s Pathogen Box [15]. Tolfenpyrad inhibits the electron transport chain complex I and impairs ATP-production [31]. The identification of the macrocyclic lactones and tolfenpyrad as potent actives in our assay validates its utility to identify anthelmintics.

**Fig. 2.**
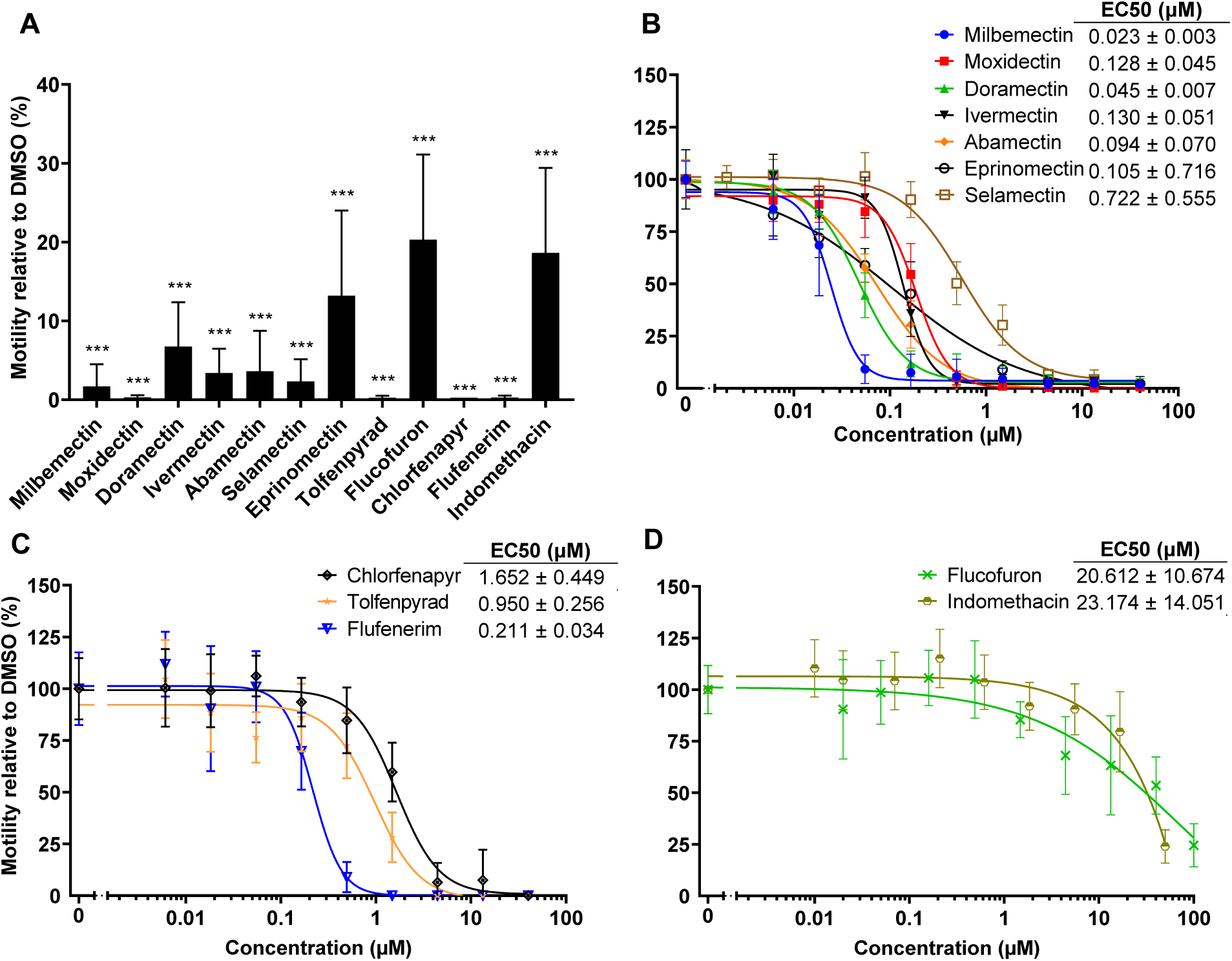
Primary and concentration-response screen data for the 12 primary hits. (**A**) Primary screening identified twelve hits in the MMV COVID and Global Health Priority Boxes. L4 were exposed to 40 µM compound and incubated for 24 h. Motility was normalized to that of the DMSO control. Data displayed represent the means ± SD values from one or two independent experiments, each performed in triplicate. Statistical analysis, comparing to the DMSO control, was conducted using a one-sample t-test: *** = p < 0.0001. (**B** – **D**) Concentration response data for the 12 primary hits. L4 were incubated with nine concentrations of compound, and the data evaluated after (**B**) 1 h (except for selamectin which was evaluated after 3 h), (**C**) 12 h or (**D**) 24 h for measuring EC_50_ values. Shown are the means ± SD values from two or three independent assays, each performed in duplicate.

Of the four remaining hits, chlorfenapyr (MMV1577458; 0.26% motility relative to control), flucofuron (MMV027339; 20.31%), flufenerim (MMV1794206; 0.26%) are pesticides, whereas indomethacin (MMV002813; 18.64%), the only hit from the COVID Box, is a non-steroidal anti-inflammatory drug (NSAID). The latter three compounds are apparently novel for their activity against *C. elegans*.

### Concentration response screens

To determine the relative potency of the 12 primary hits, concentration response curves were generated. Also evaluated was the time-to-effect, whereby compounds were classified by the time point at which an EC_50_ value was first calculable, namely, after 1 h (**Fig. 2B**), 12 h (**Fig. 2C**) or 24 h (**Fig. 2D**). All macrocyclic lactones, except selamectin, demonstrated EC_50_ values <1 µM after 1 h. The current EC_50_ value of 0.130 ± 0.051 µM for ivermectin is similar to the value of 0.19 ± 0.01 µM previously recorded after 90 min [13]. Selamectin was somewhat slower acting with an EC_50_ value after 3 h of 0.722 ± 0.555 µM. After 12 h, the EC_50_ values for chlorfenapyr, tolfenpyrad and flufenerim were 1.652 ± 0.449 µM, 0.950 ± 0.256 µM and 0.211 ± 0.034 µM, respectively (**Fig. 2C**). The value measured here for tolfenpyrad after 12 h was higher than that previously recorded after 24 h (0.20 ± 0.04 µM) [15], but still within one order of magnitude. After 24 h, the EC_50_ values for the remaining two hits, flucofuron and indomethacin, were 20.612 ± 10.674 µM and 23.174 ± 14.051 µM, respectively (**Fig. 2D**).

### HEK293 cell cytotoxicity assay

The concentration-dependent cytotoxicity of the 12 hit compounds was measured using HEK293 cells after 48 h (**Fig. 3**). The macrocyclic lactones and tolfenpyrad were relatively non-toxic with CC_50_ values between 10 and 34 µM (**Fig. 3A**). Likewise, chlorfenapyr, flufenerim and indomethacin were essentially non-toxic, with CC_50_ values between 21 and >100 µM. More potent cytotoxicity was measured for flucofuron (0.453 ± 0.331 µM; **Fig. 3B**).

**Fig. 3.**
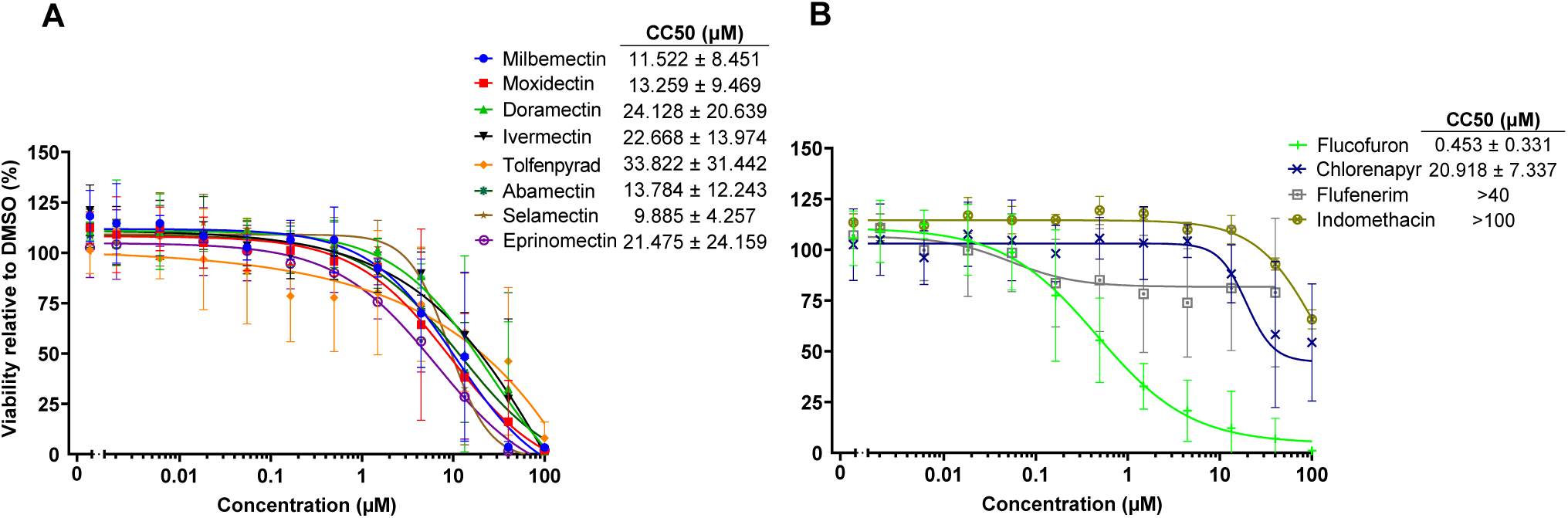
Cytotoxicity of the 12 hit compounds. HEK293 cell viability was measured after 48 h for the seven macrocyclic lactones and tolfenpyrad (**A**), and for flucofuron, chlorfenapyr, flufenerim and indomethacin (**B**). Data represent the means ± SD from two experiments each performed in duplicate

## Discussion

Parasitic nematode infections are a global health problem and a continual challenge to agricultural productivity [1,2] due to the emergence and establishment of drug resistance. Thus, new drugs are needed. In this context, we screened the MMV’s recently-released COVID and GHP Boxes against the free-living nematode, *C. elegans,* which has a recognized utility in the identification of anthelmintics [12–15].

We employed the WMicroTracker ONE infrared-based motility device which we preliminarily reported as a useful screening tool [32]. The device has also been employed with various nematode species for drug repurposing and to identify active natural products [33, 34]. It is economical, simple to use and its associated software is easy to learn. Also, the quantitative outputs allow for comparisons of compound efficacy, a notable alternative to manually estimating mortality (*e.g*., [12, 35]), which can be time-consuming and require an understanding of nematode morphology.

After optimizing the assay for worm numbers, DMSO concentration and assay volume, we performed single concentration screens. Among the potent hit compounds were seven established macrocyclic lactone anthelmintics, which were, in the main, fast-acting with sub-micromolar EC_50_ values after 1 h.

Two other hits, tolfenpyrad and chlorfenapyr, were identified with EC_50_ values measurable after 12 h. Of these, chlorfenapyr, a pyrrole class pro-insecticide, was recently employed as a test compound in a screen to measure the response variations among eight *C. elegans* strains to environmental toxicants [36]. In animals, dealkylation of an ethoxymethyl group on chlorfenapyr, reveals the active metabolite, tralopyril, which uncouples mitochondrial oxidative phosphorylation to disrupt ATP-production leading to death [37]. Although relatively non-toxic to HEK293 cells (EC_50_ = 20.918 ± 7.337 µM), there have been poisoning cases with chlorfenapyr [38, 39, 40], likely undermining its direct repurposing potential as an anthelmintic.

The screen also identified three novel anthelmintics, flufenerim, flucofuron and indomethacin. An EC_50_ value was calculable for flufenerim after 12 h, whereas the values for flucofuron and indomethacin were only calculable after 24 h. The pyrimidineamine, flufenerim, is a potent plant insecticide [41]. Although its mode of action is not confirmed, it inhibits acetylcholinesterase *in vitro* and *in vivo* [41]. Acetylcholinesterase is an established anthelmintic/insecticide drug target [42, 43, 44] and, if the primary target here, then it may be possible to modify the compound’s structure to enhance specificity and selectivity.

Flucofuron is a bisarylurea pesticide that has been used to mothproof wool [45]. It is a relatively modest inhibitor of worm motility (EC_50_ = 20.612 ± 10.674 µM) with some HEK293 cell toxicity (CC_50_ = 0.453 ± 0.331 µM). In cancer cells, flucofuron and its lipophilic, electron-withdrawing analogues, uncouple mitochondria via the fatty acid-activated mechanism, thereby disrupting ATP-production to cause apoptosis [46]. Disrupting oxidative phosphorylation is a key anti-parasitic strategy [12, 47, 48], and like flufenerim, flucofuron may be chemically modifiable to generate selectivity and specificity to the nematode target.

The third novel hit, the indole-based NSAID, indomethacin, was not cytotoxic to HEK293 cells. Like other NSAIDs, indomethacin inhibits cyclooxygenase (COX) enzymes and their production of prostaglandins, which mediate inflammation [49]. Interestingly, no COX-like enzyme is present in *C. elegans* [50, 51, 52], suggesting that indomethacin may act on a novel target(s). COX-2 selective indomethacin analogs with excellent safety profiles have been synthesized [53]. Thus, the opportunity exists to possibly improve on the modest potency demonstrated here for indomethacin *vs. C. elegans* by further screening these or other analogs, combined with genetic approaches to identify the target(s).

## Limitation

Although *C. elegans* is a proven model for anthelmintic drug discovery [12–15], the bioactives discovered here would need to be confirmed in screens of parasitic nematodes.

## Abbreviations

DMEM: Dulbecco’s Modified Eagle Medium
DMSO: Dimethyl sulfoxide
EDTA: Ethylene diamine tetra acetic acid
FBS: Fetal bovine serum
GHP: Global Health Priority
HEK: human embryonic kidney
MMV: Medicines for Malaria Venture

## Ethics approval

None to declare.

## Consent for publication

Not applicable.

## Acknowledgements

The authors thank Karol R. Francisco, Bobby Lucero, Thaiz Rodrigues Teixeira and Dilini K. Amarasinghe for their generous advice regarding the data analysis and revision of the manuscript. The authors also thank Sreekanth Chalasani of the Salk Institute for Biological Sciences, La Jolla, CA, and Emily Troemel of Department of Cell and Developmental Biology, School of Biological Sciences, University of California San Diego, CA, for providing the N2 strain of *C. elegans* and the OP50 strain of *Escherichia coli*, respectively.

## Authors’ contributions

All authors conceptualized the project together. YUS and LJL designed and optimized the *C. elegans* screening assay. YUS screened the MMV compounds with both *C. elegans* and the HEK293 cells, and performed the data analyses. All authors contributed to the writing and approval of the final manuscript.

## Availability of data and materials

All data supporting the results of this article are included in this article and its additional files. Any materials or databases generated in this study are available from the corresponding author upon reasonable request.

## Additional material

**Additional Table 1.**
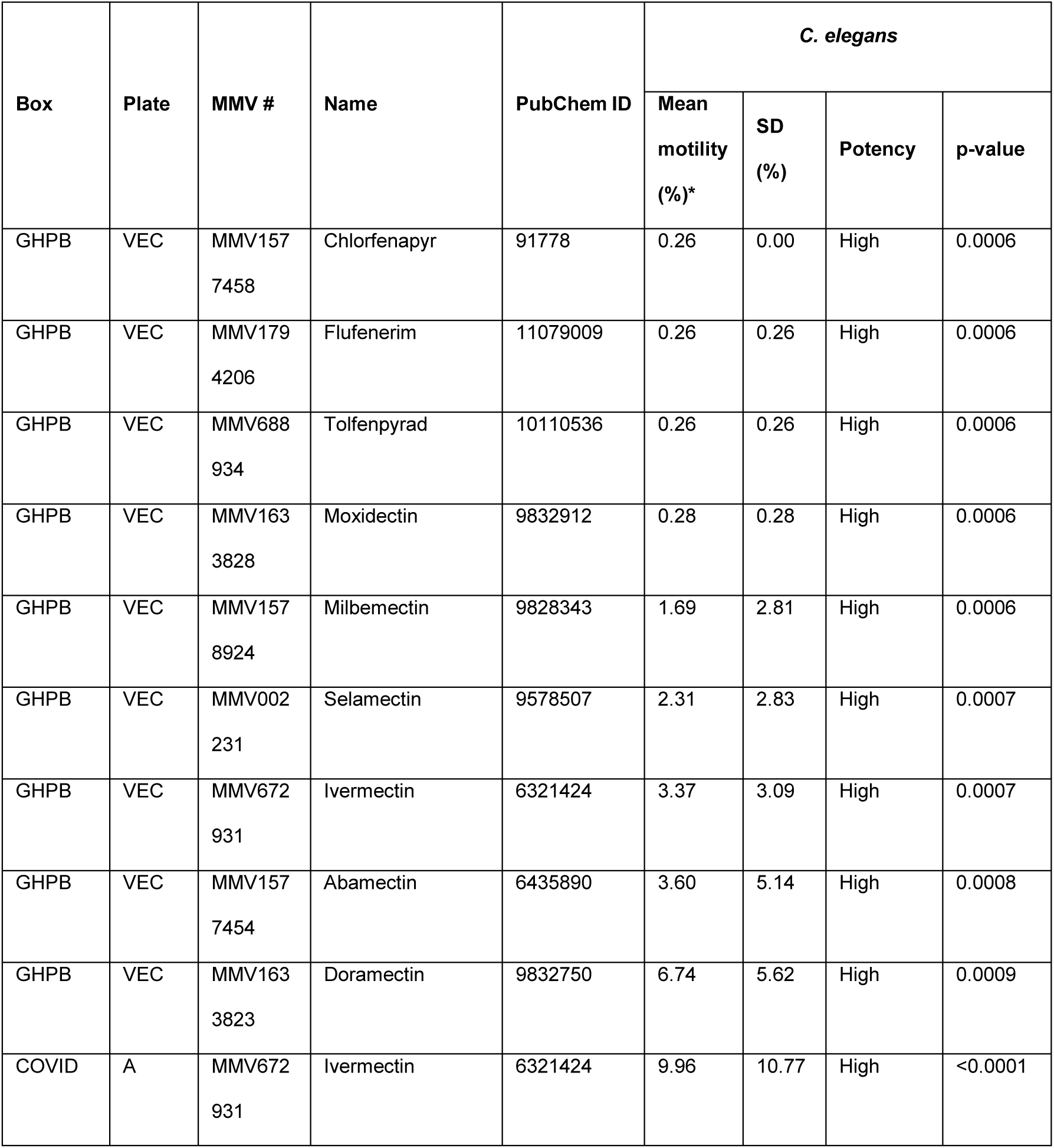
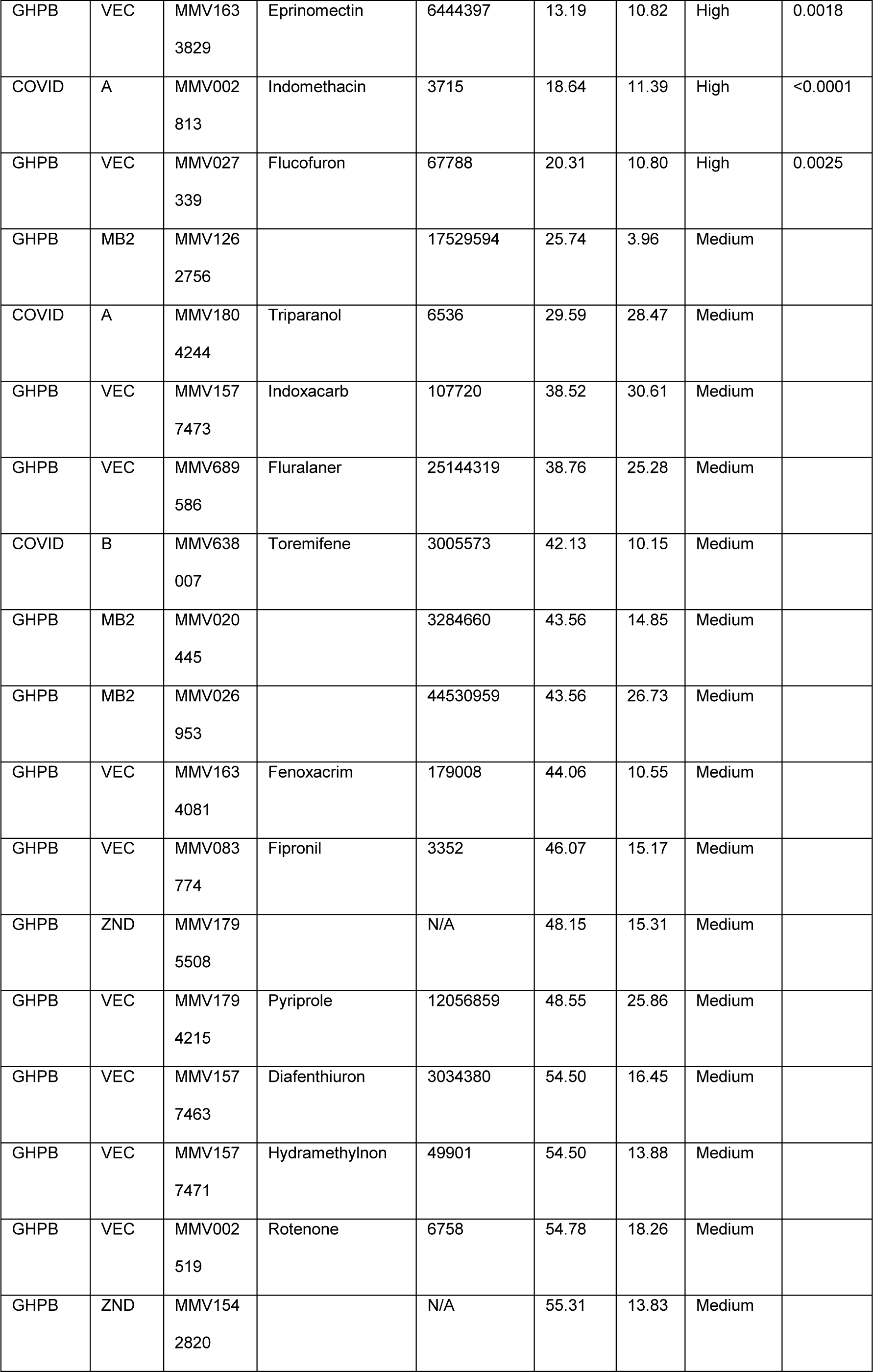
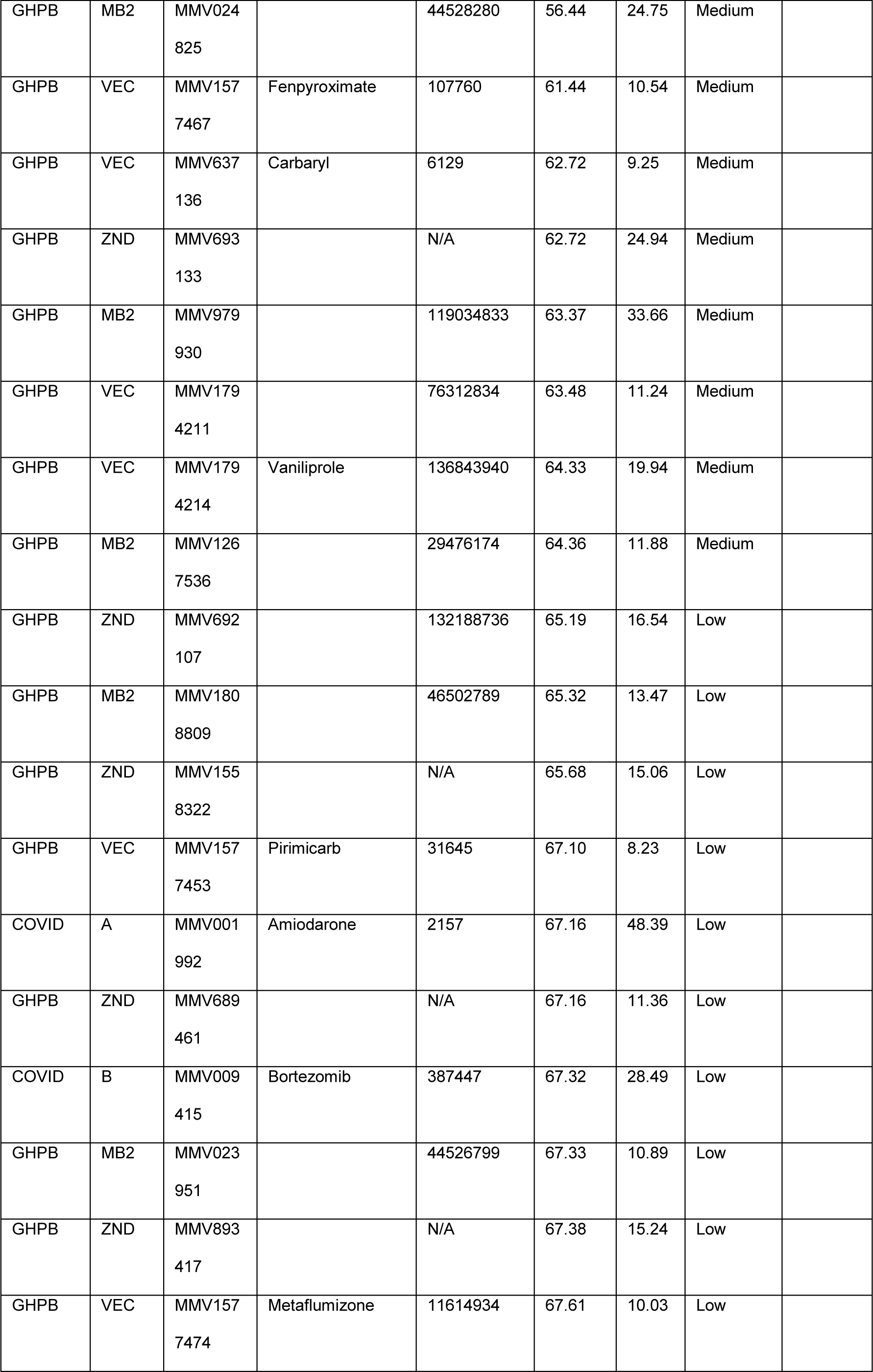
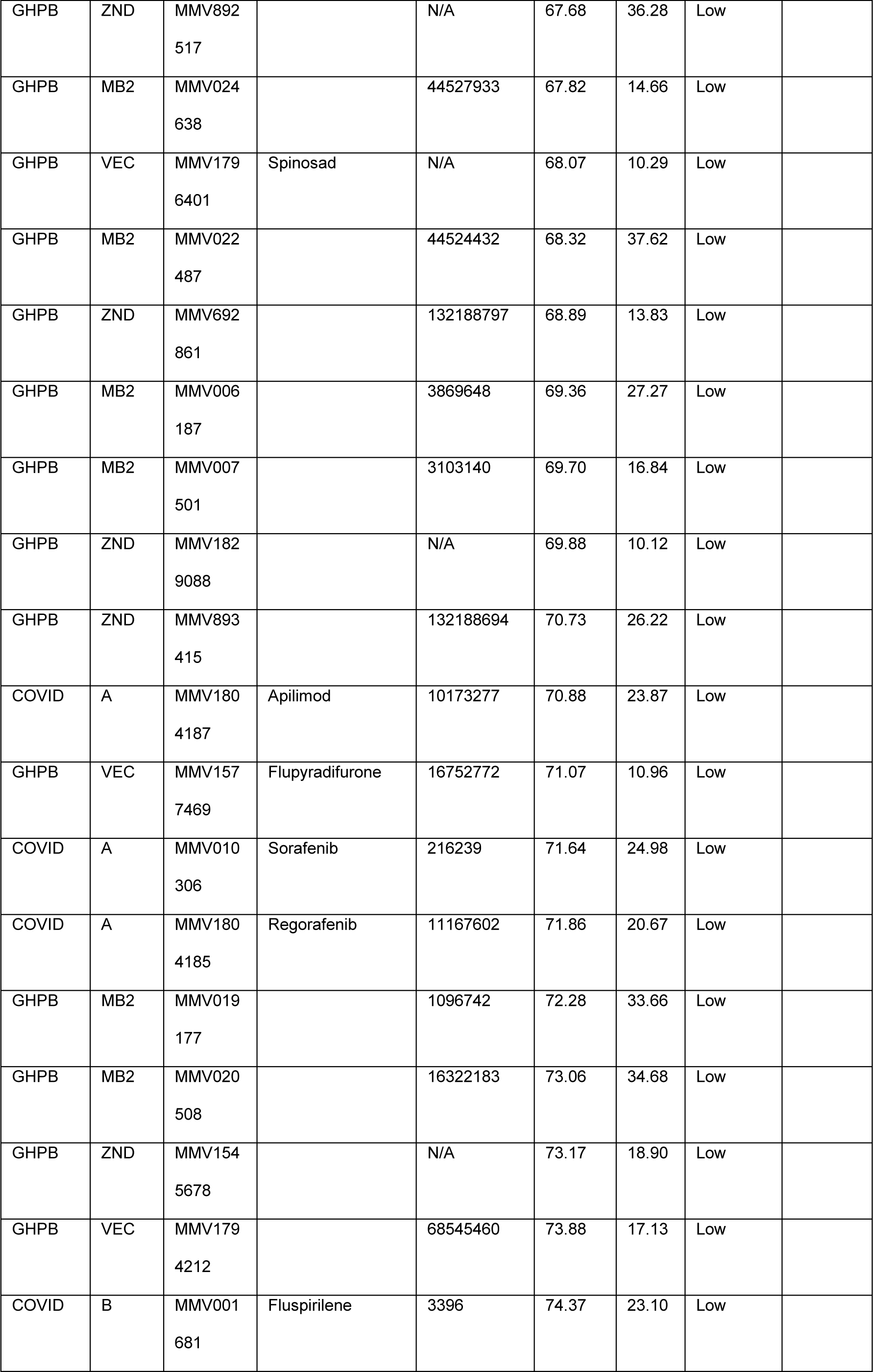
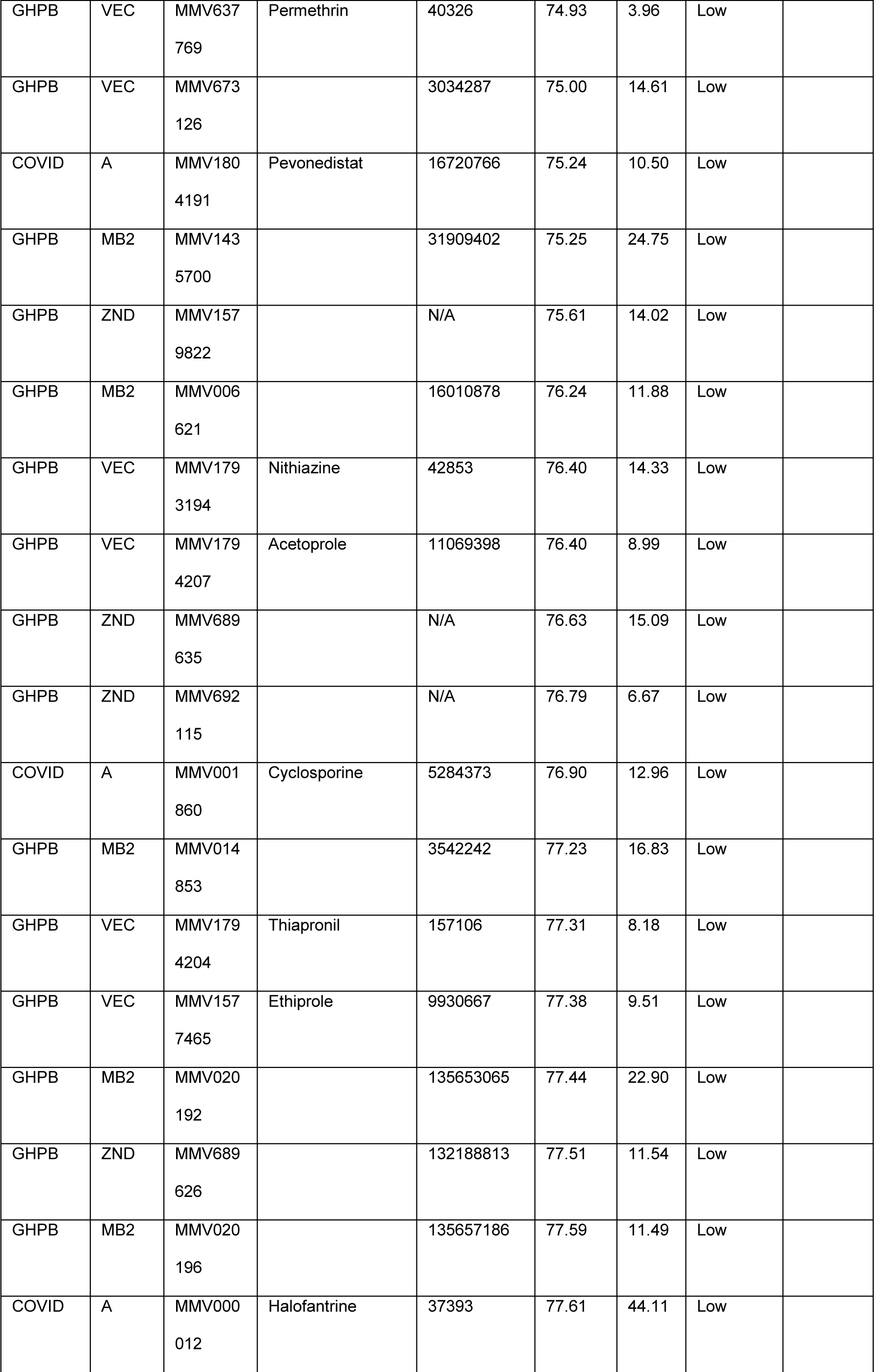
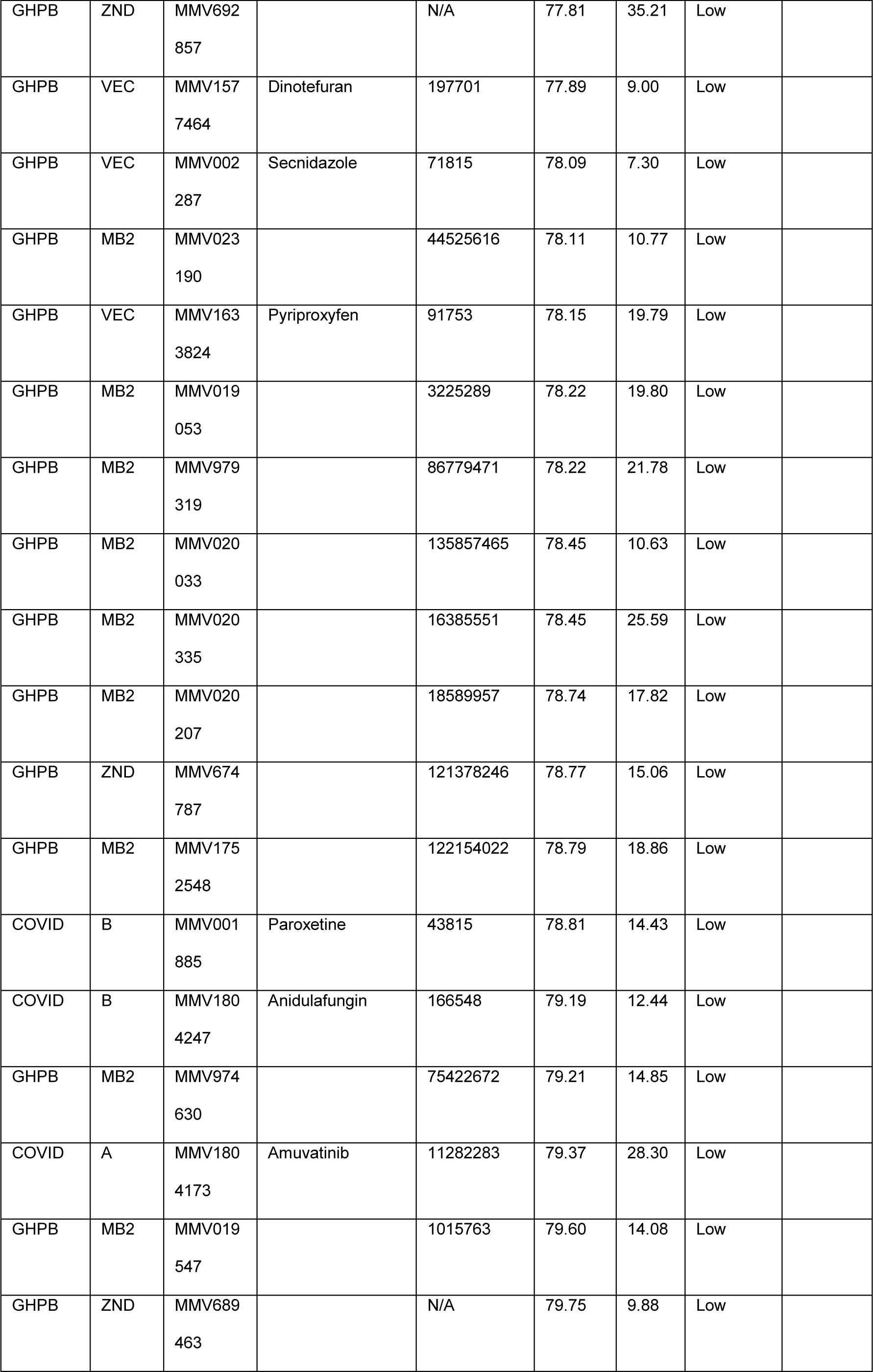
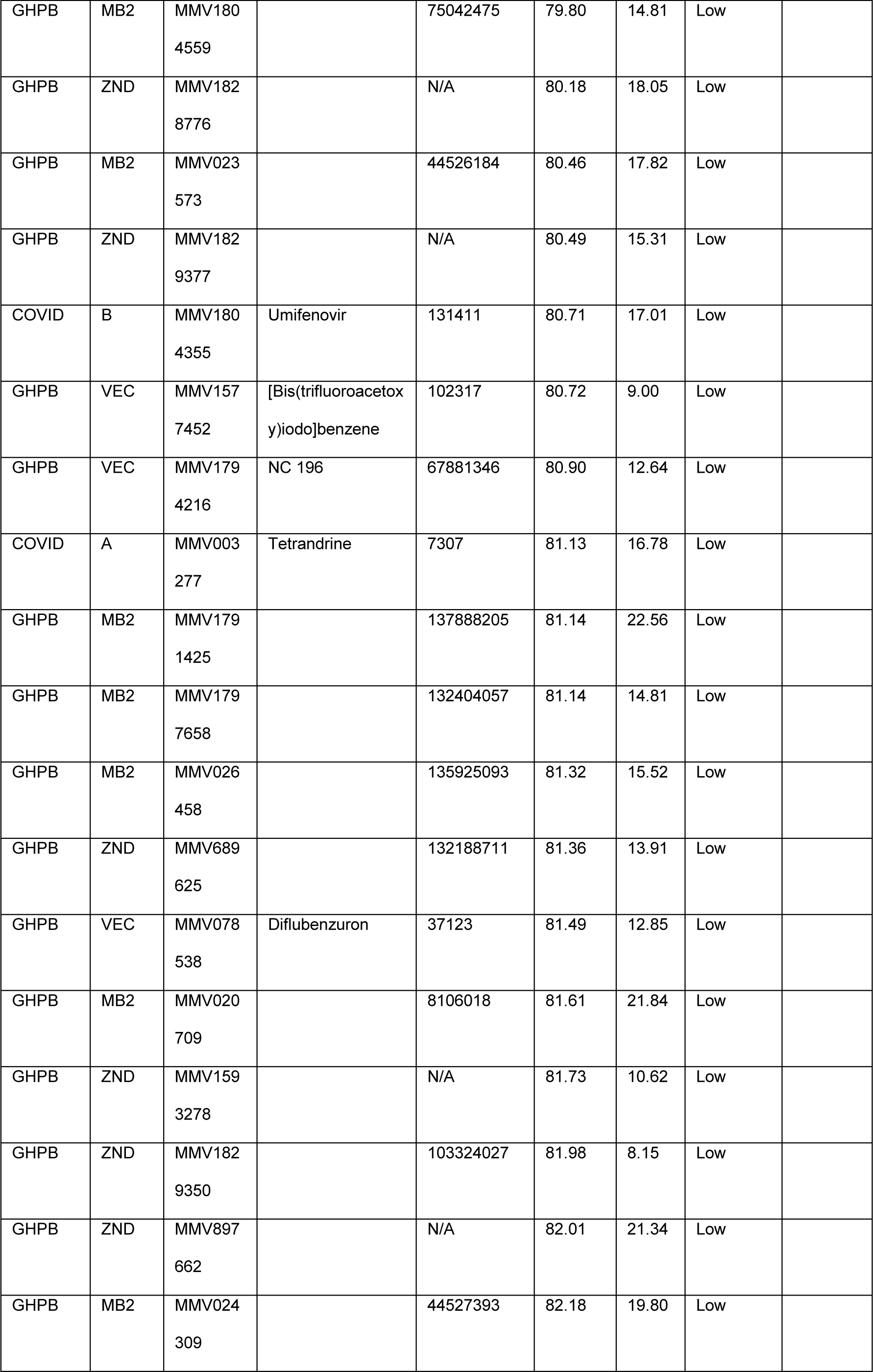
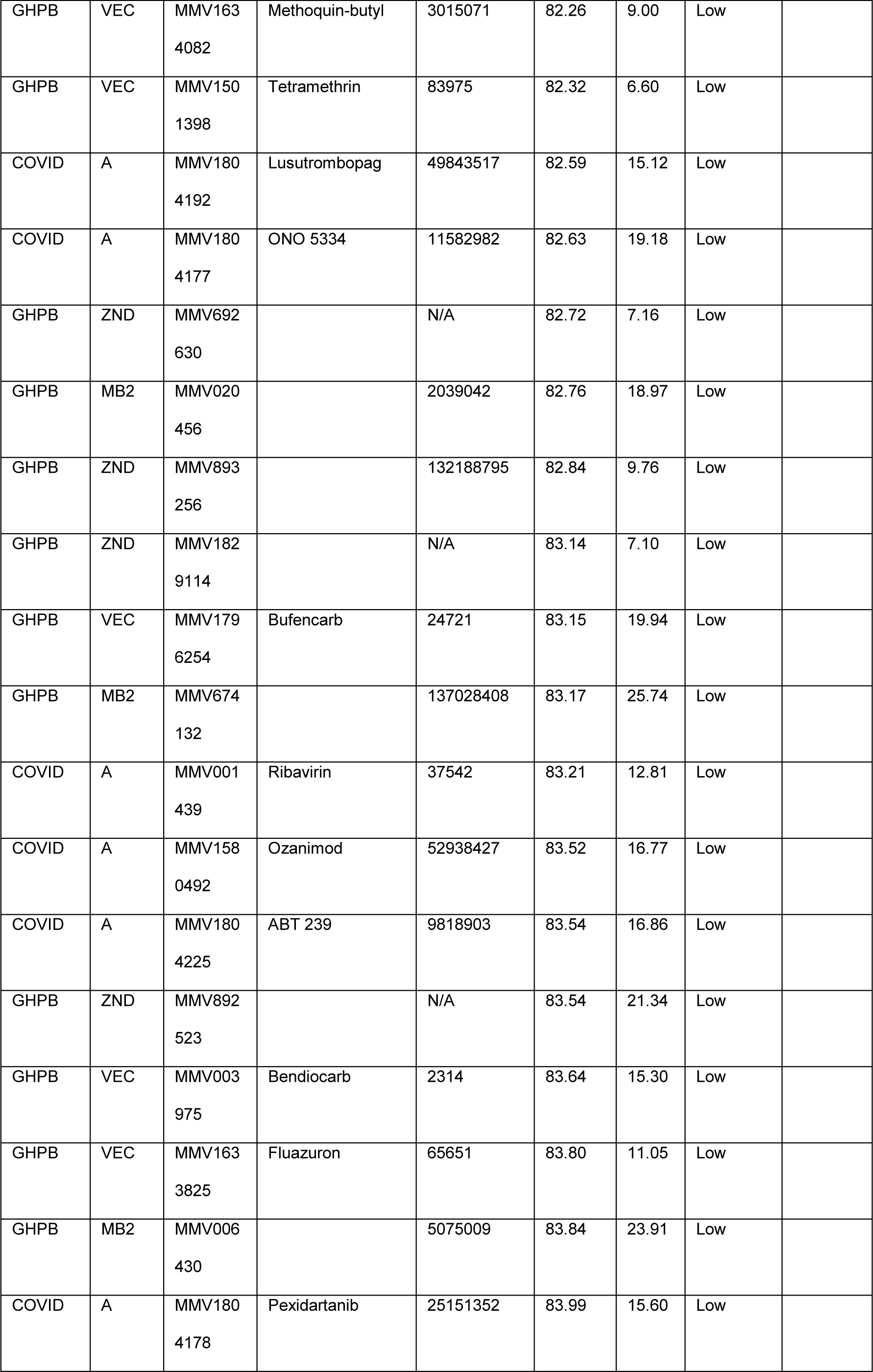
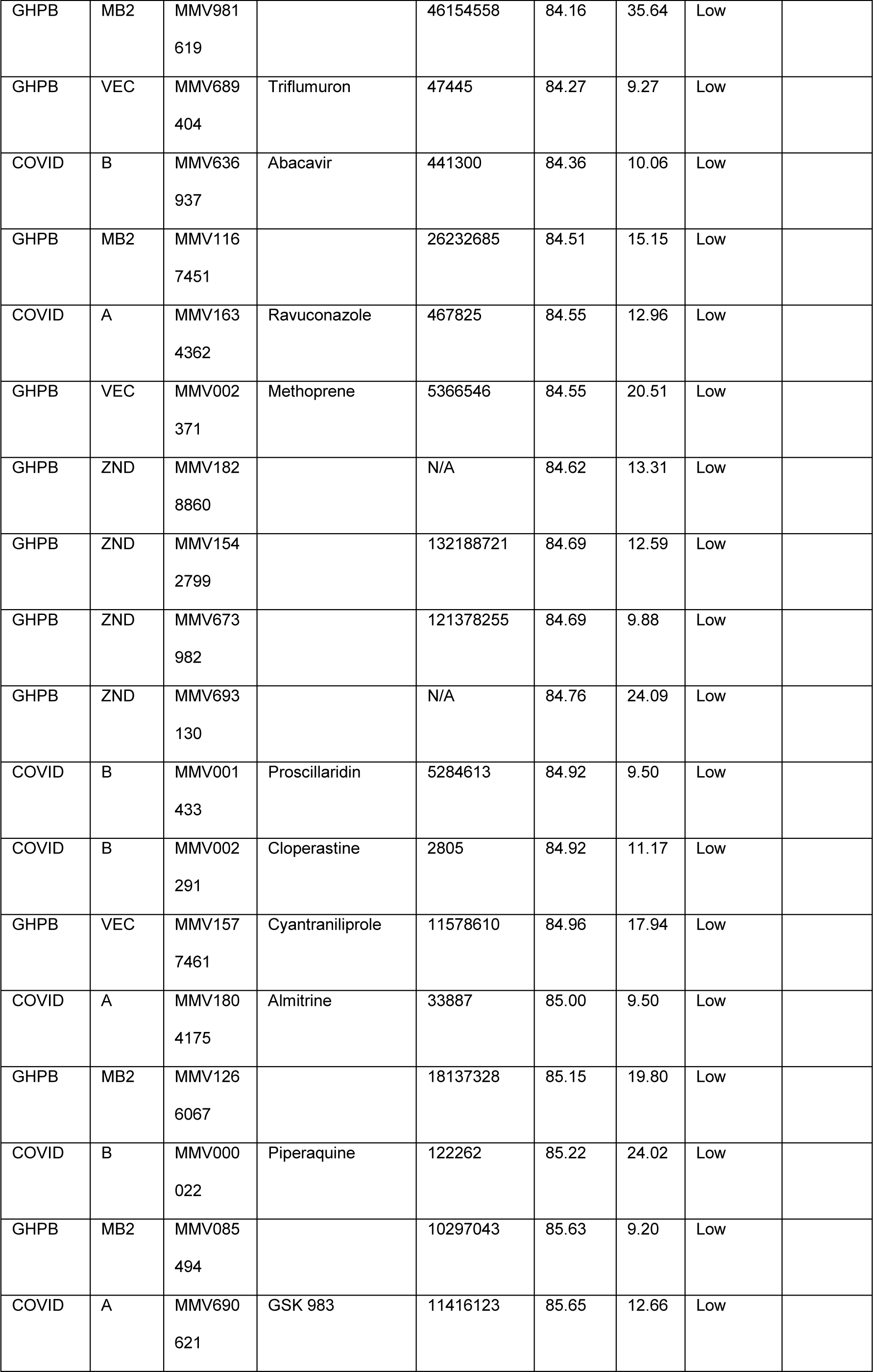
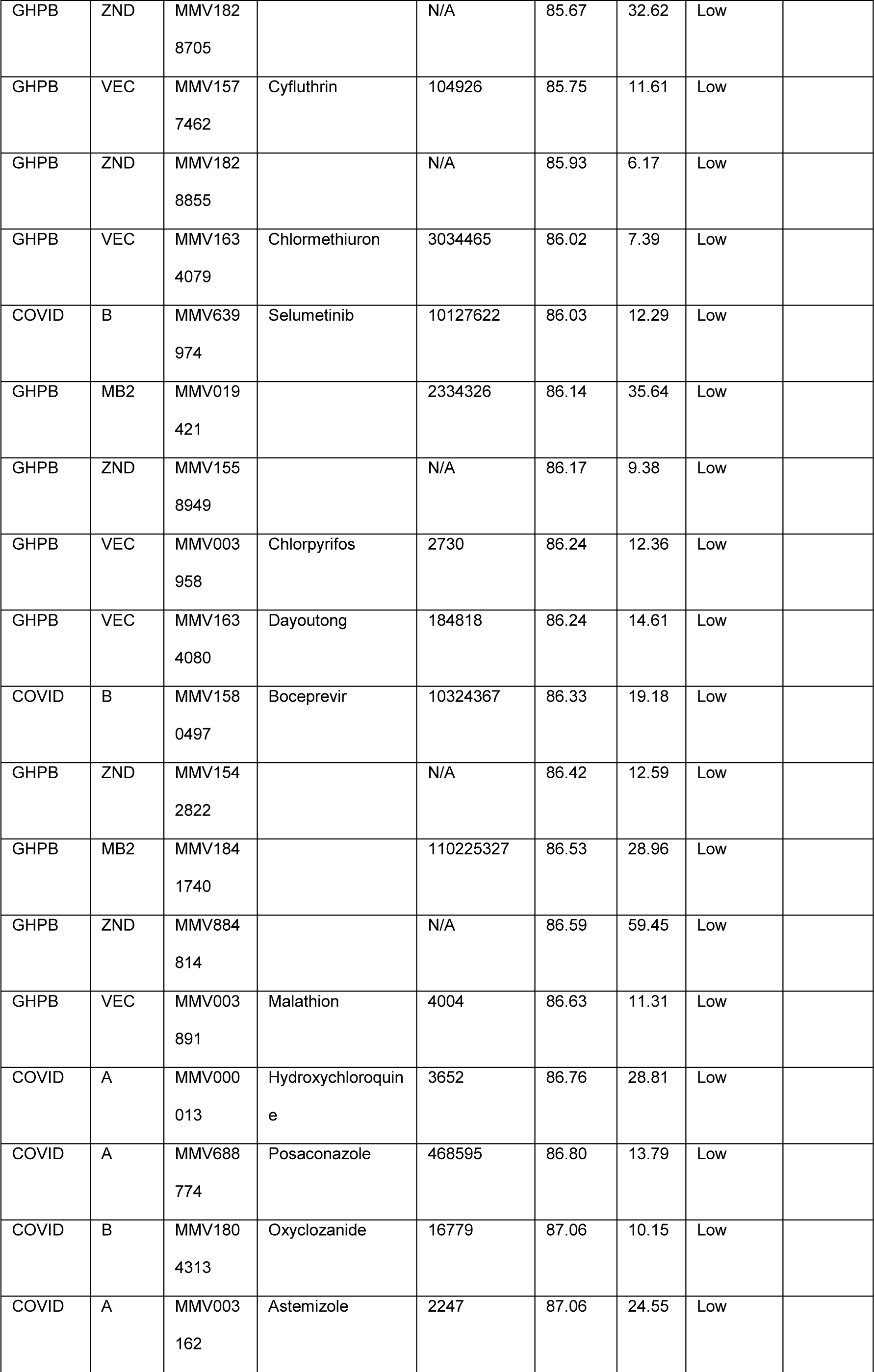
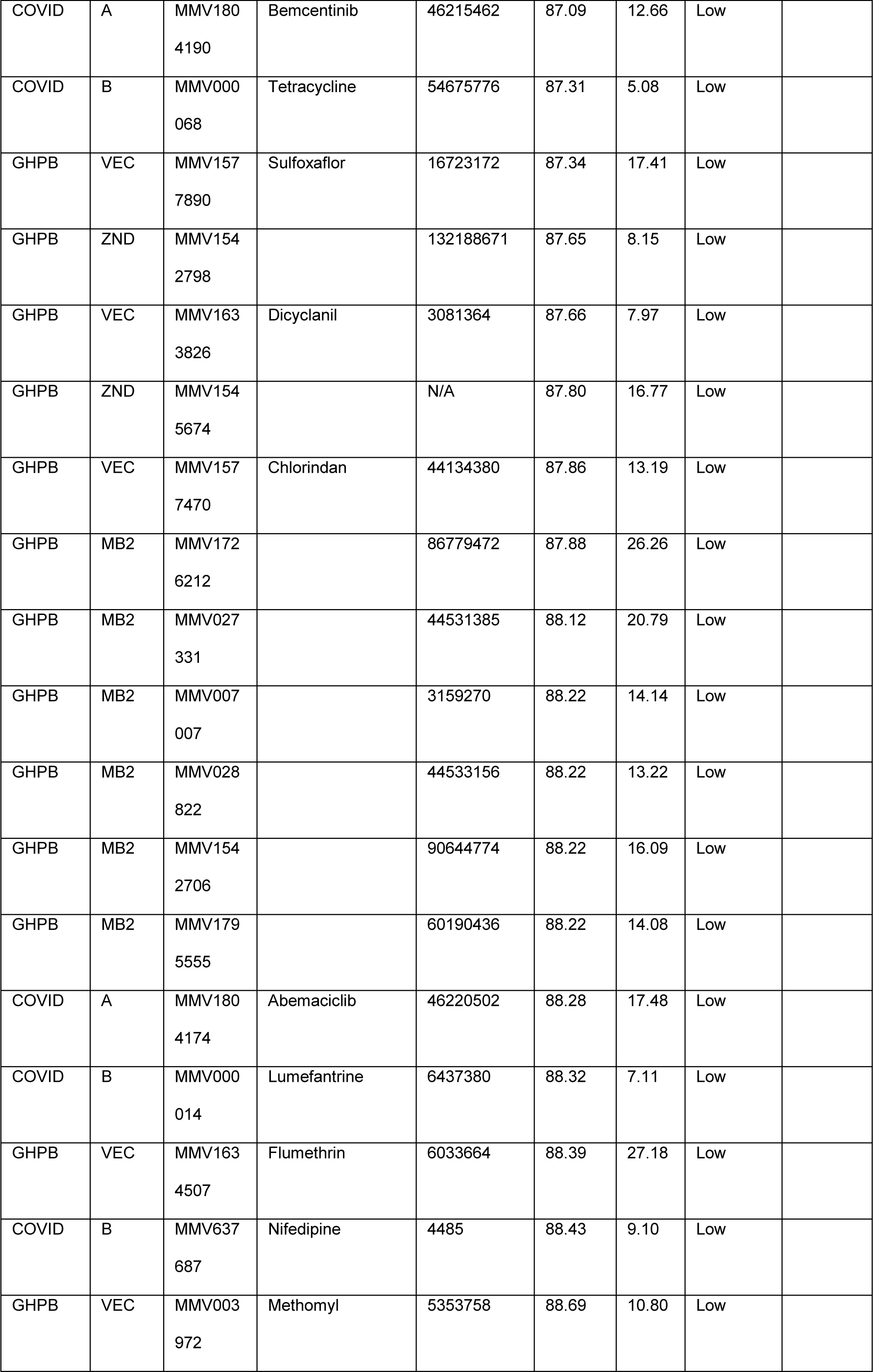
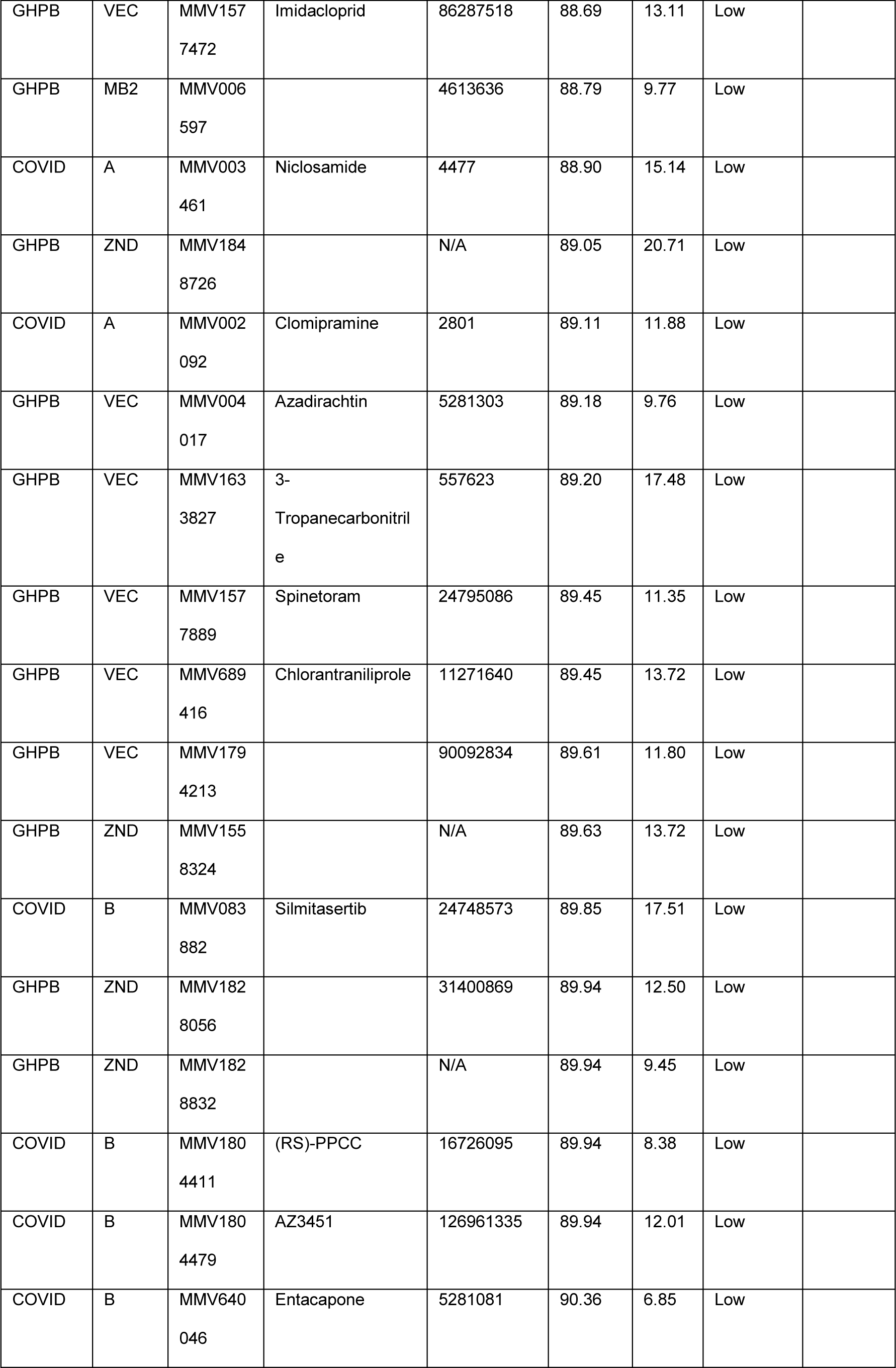
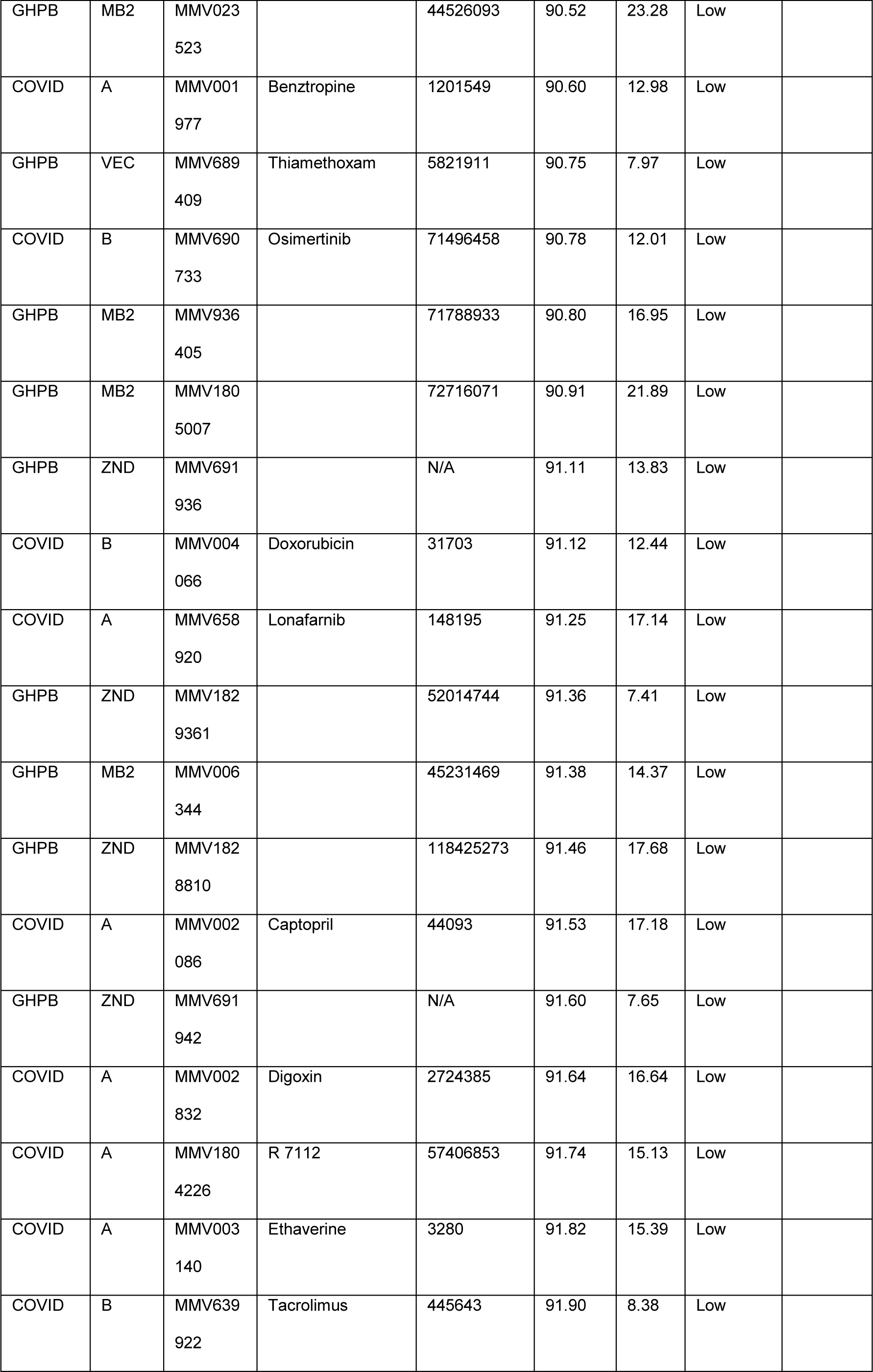
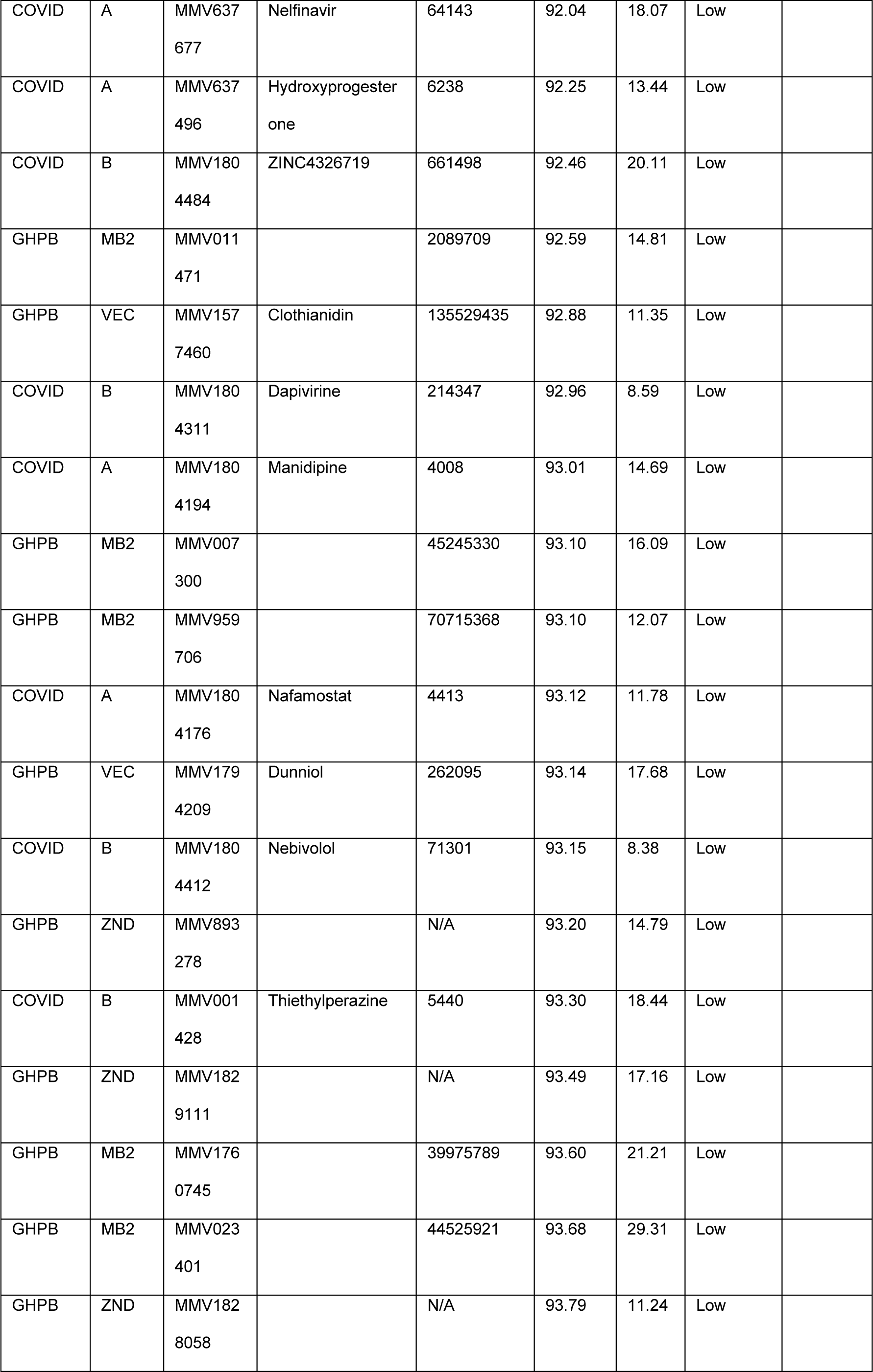
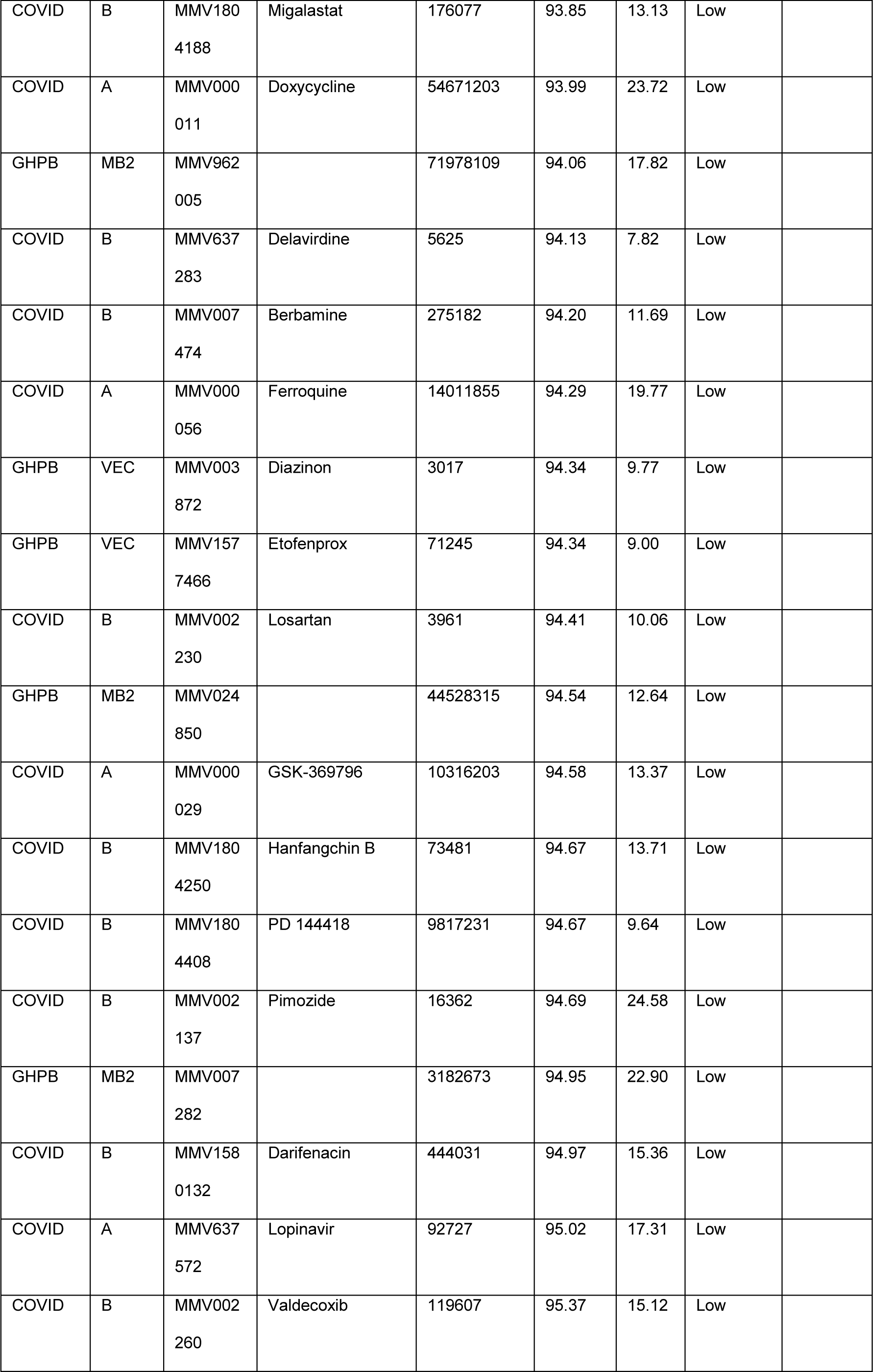
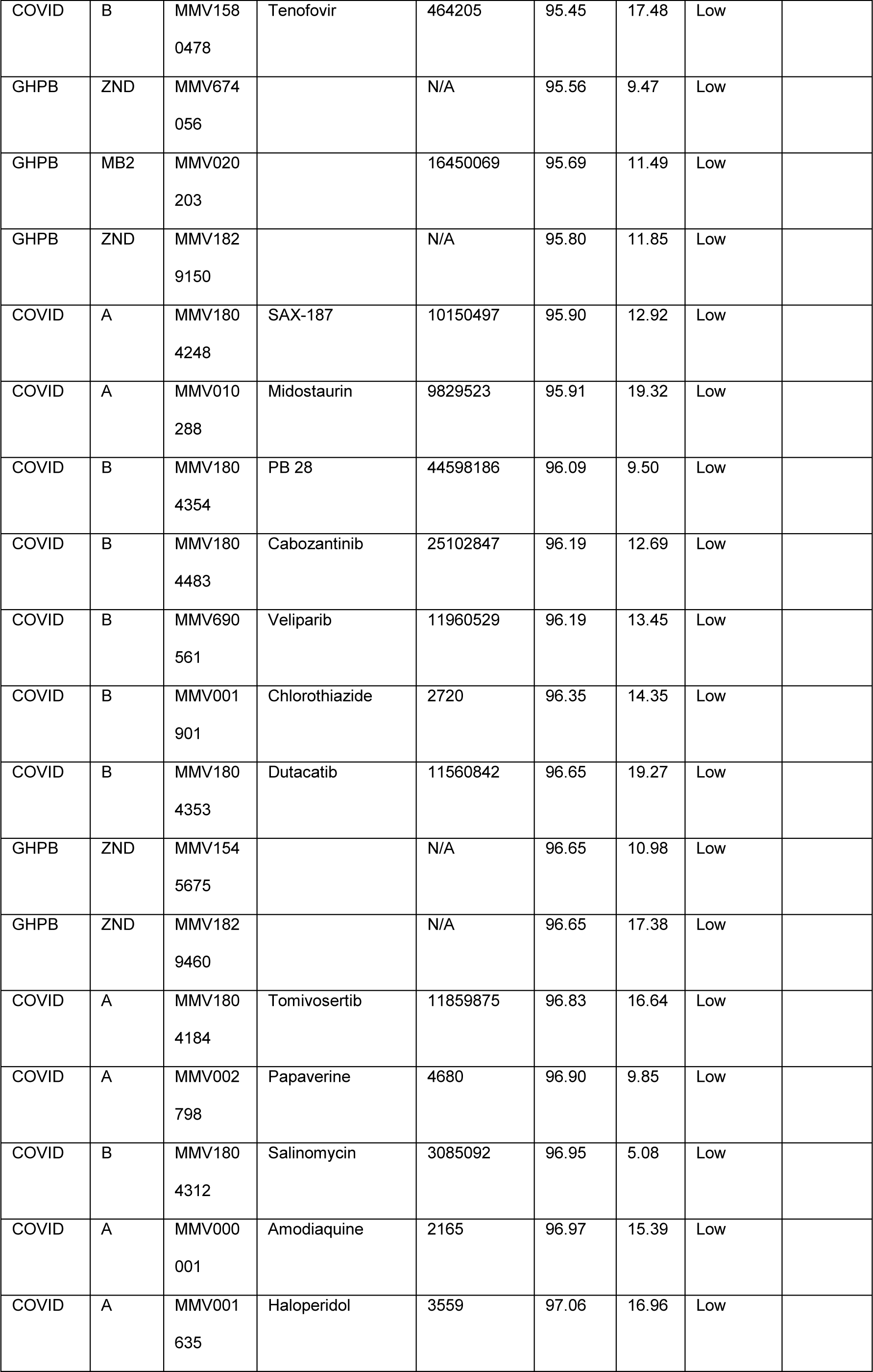
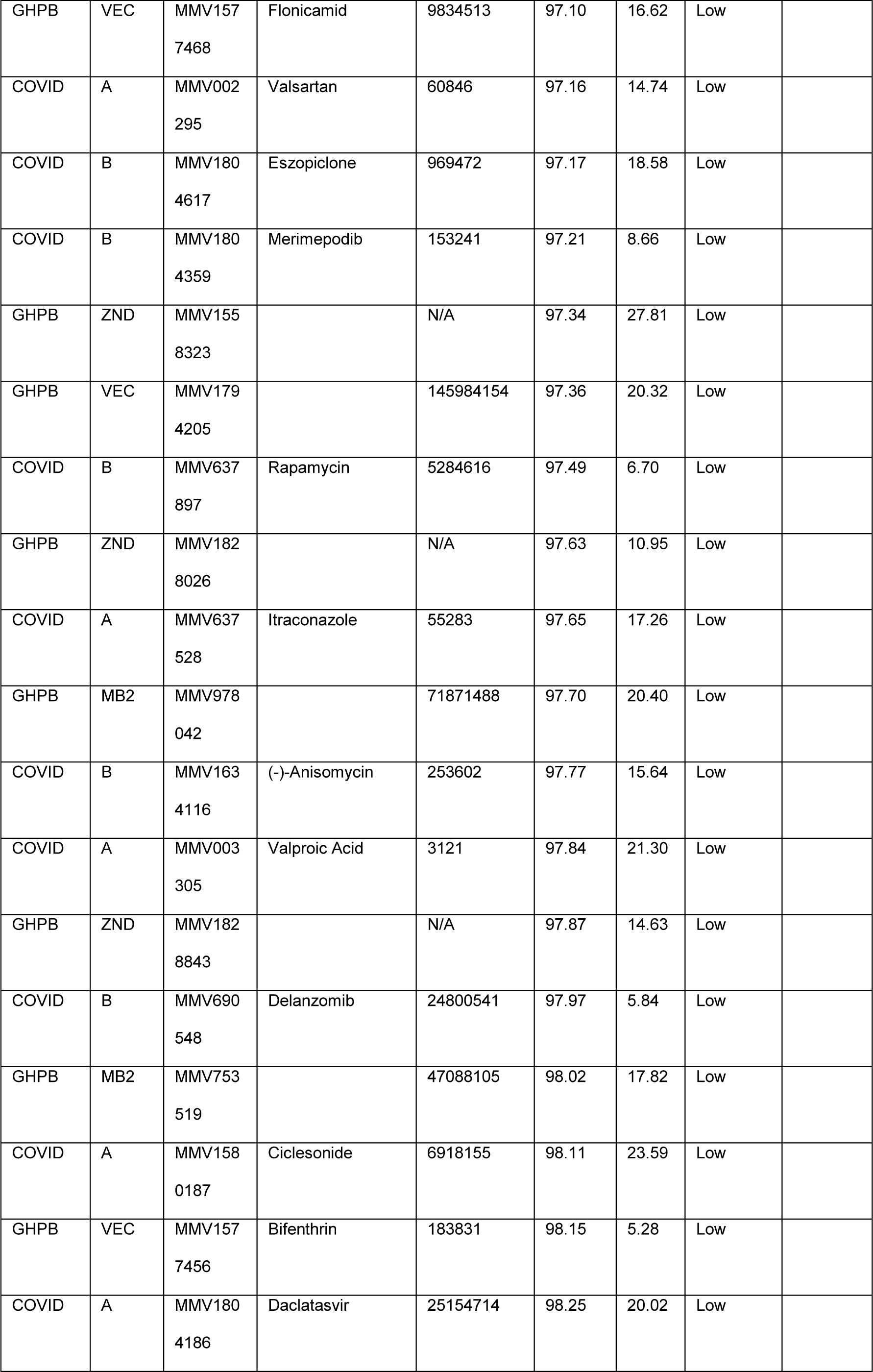
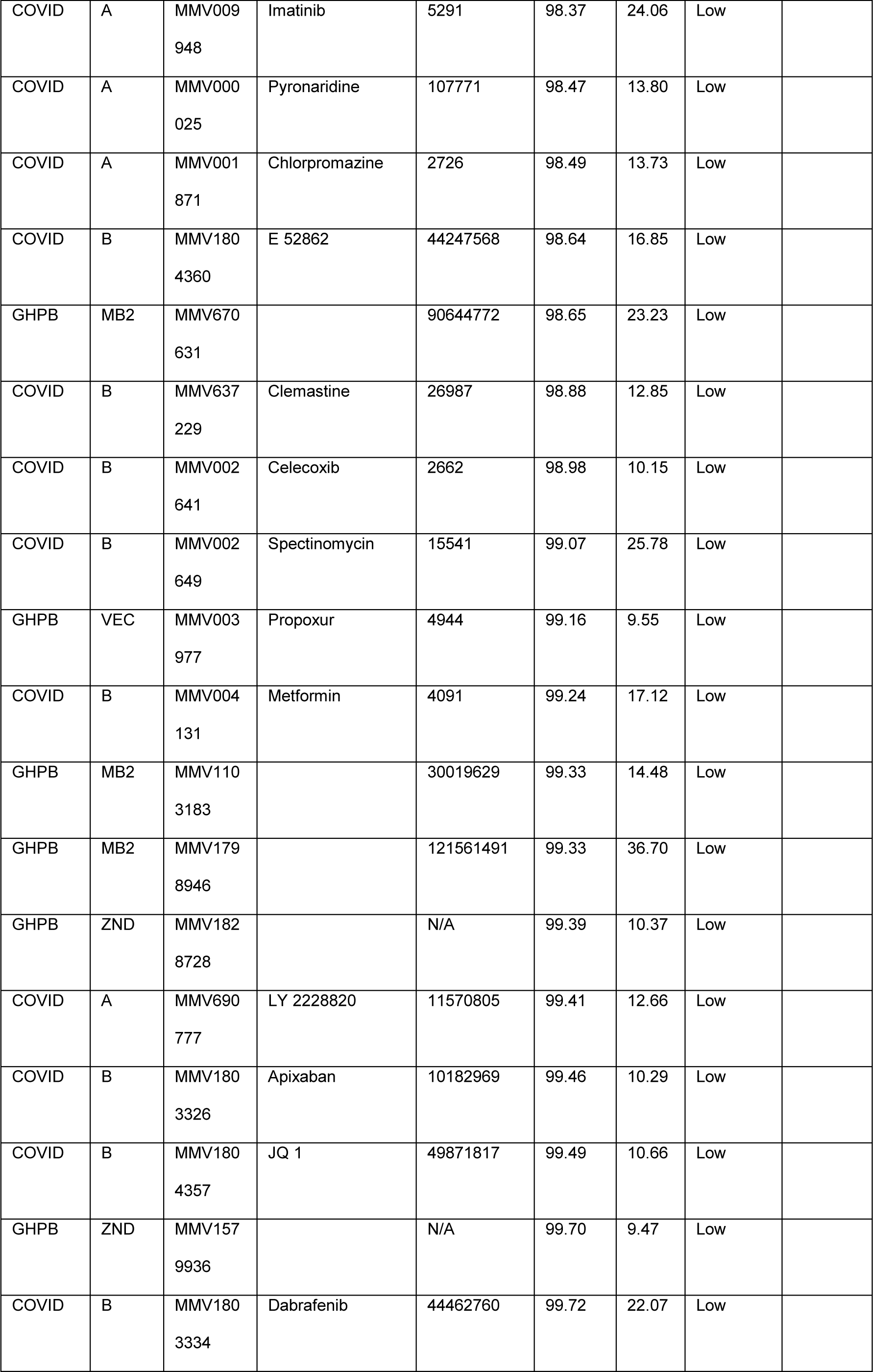
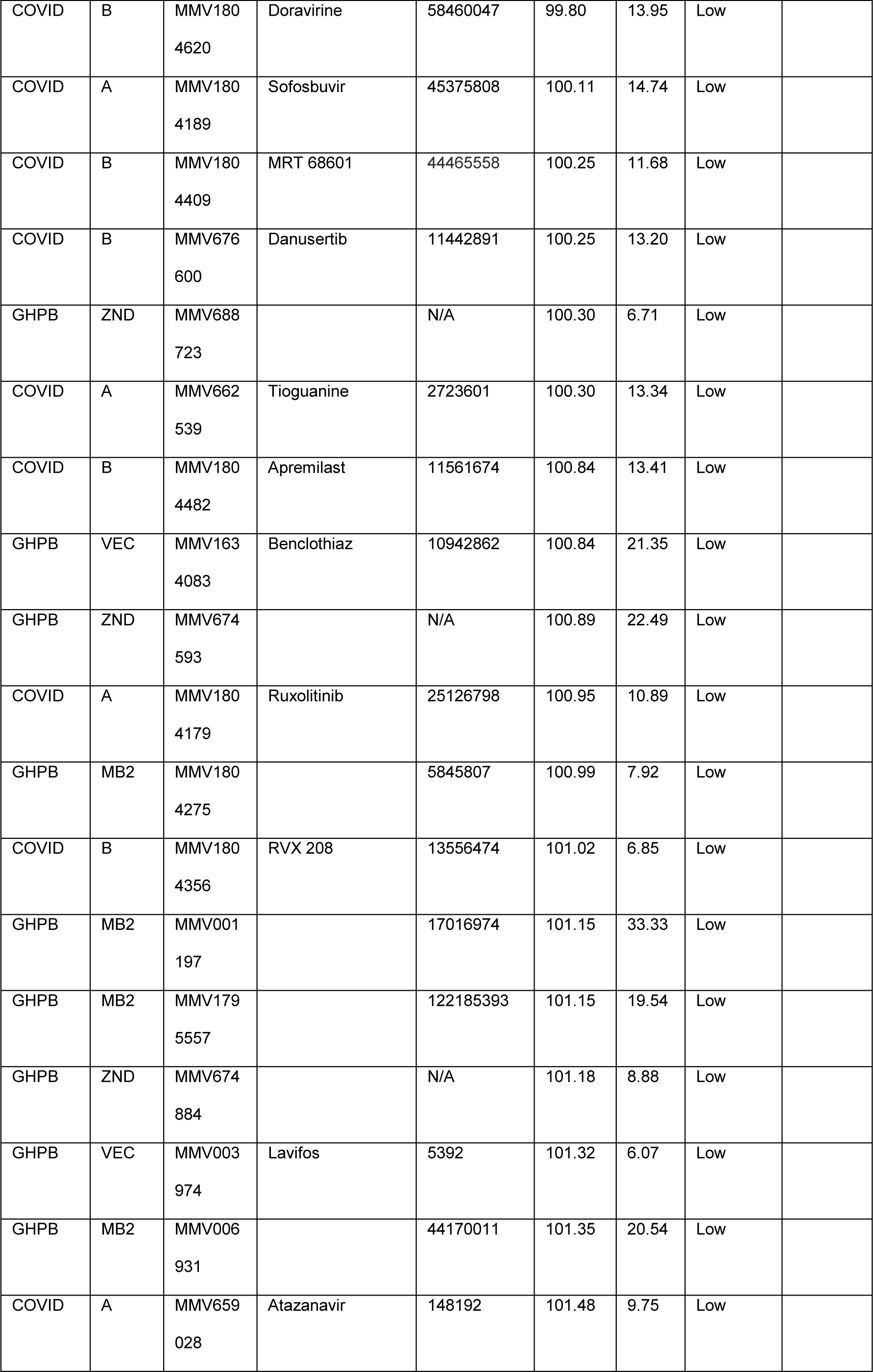
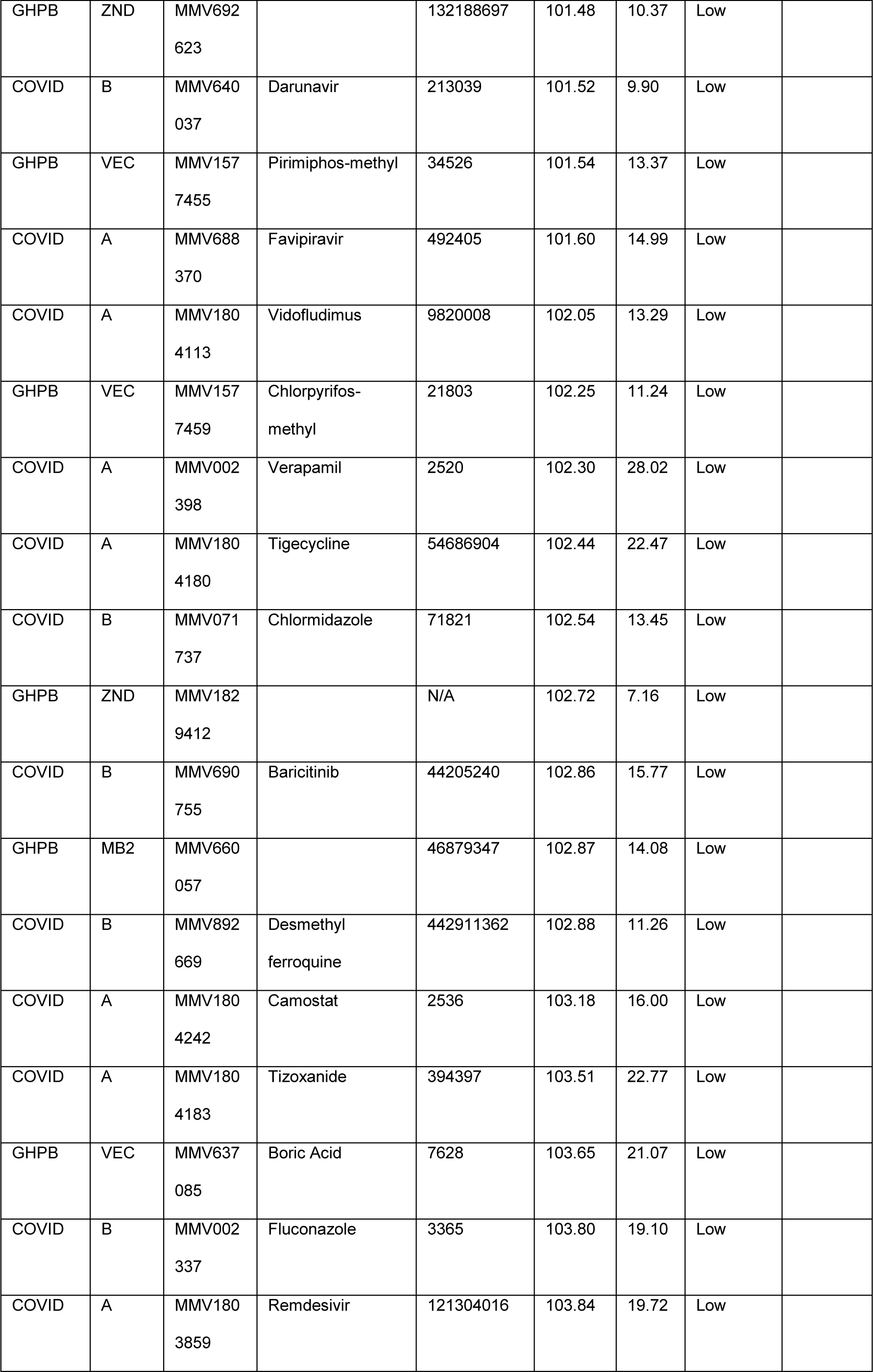
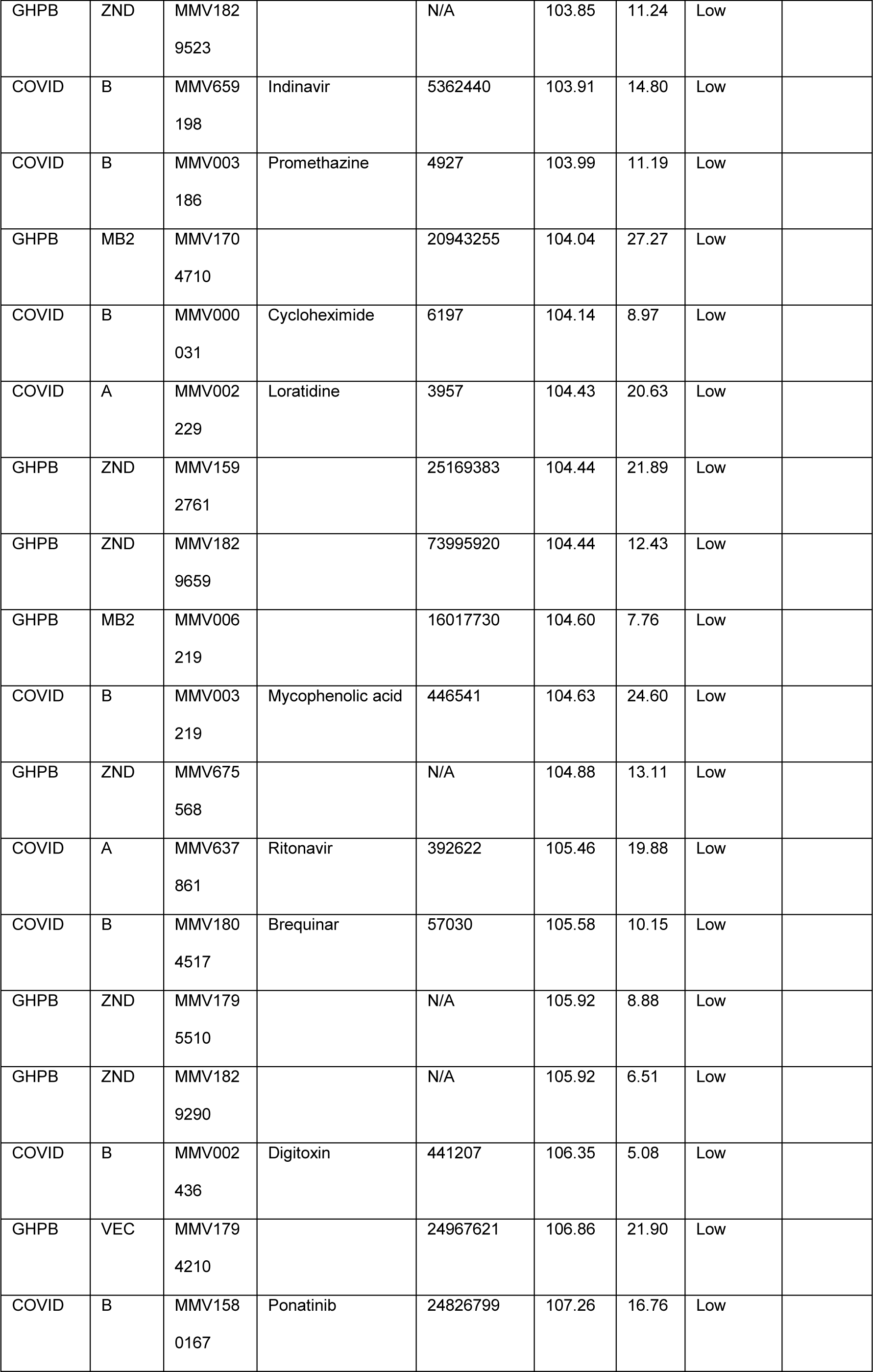
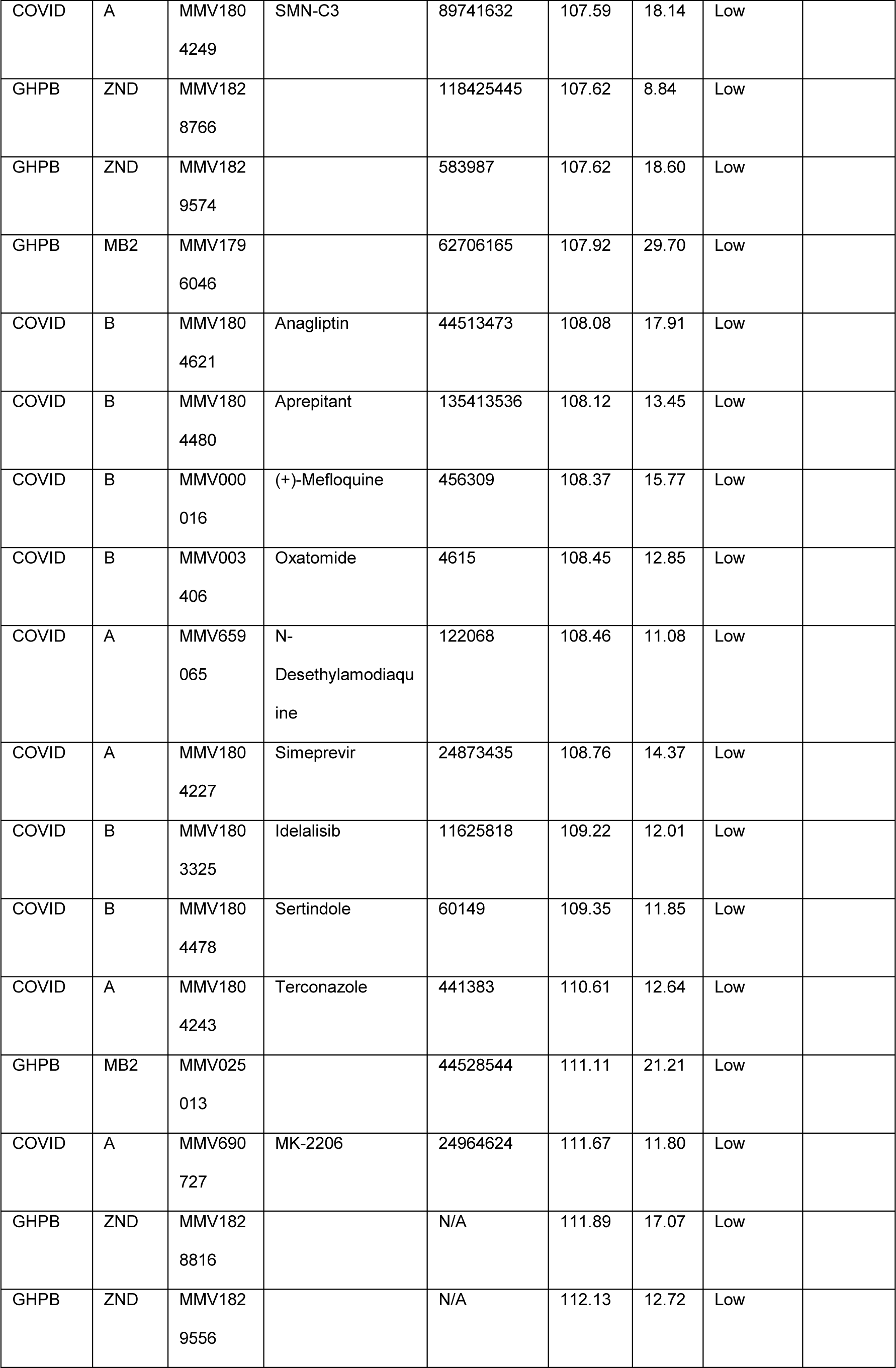
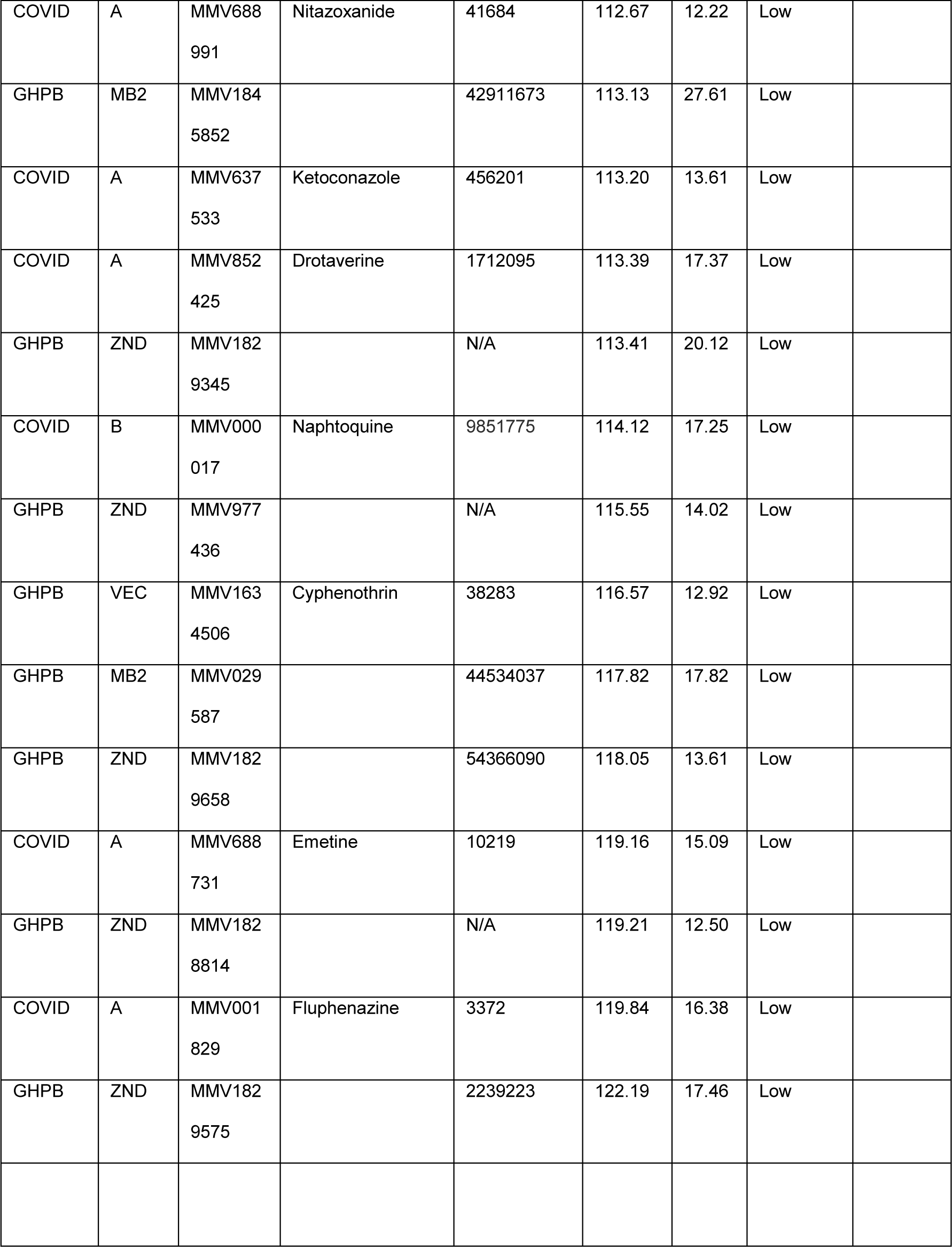
Primary screen activities of the 400 compounds in the MMV COVID and Global Health Priority Boxes. Compounds were tested at 40 µM for modulation of *C. elegans* motility after 24 h. Relative to DMSO control, the degree to which worm motility was decreased was characterized as potent <25%, moderate ≥25% but <65%, or low = ≥65%. One (Global health priority box (GHPB)) or two assays (COVID box) in triplicate was performed. P-values were calculated using a one-sample *t*-test for potent compounds only.

## References

1. Mendoza-de Gives P. Soil-borne nematodes: impact in agriculture and livestock and sustainable strategies of prevention and control with special reference to the use of nematode natural enemies. pathogens. 2022; doi:10.3390/pathogens11060640.

2. Montresor A, Mupfasoni D, Mikhailov A, Mwinzi P, Lucianez A, Jamsheed M, Gasimov E, Warusavithana S, Yajima A, Bisoffi Z, Buonfrate D, Steinmann P, Utzinger J, Levecke B, Vlaminck J, Cools P, Vercruysse J, Cringoli G, Rinaldi L, Blouin B, Gyorkos TW. The global progress of soil-transmitted helminthiases control in 2020 and World Health Organization targets for 2030. PLoS Negl Trop Dis. 2020; doi:10.1371/journal.pntd.0008505.

3. Keiser J, Utzinger J. The drugs we have and the drugs we need against major helminth infections. Adv Parasitol. 2010;73:197–230.

4. Holden-Dye L, Walker RJ. Anthelmintic drugs and nematicides: studies in *Caenorhabditis elegans*. WormBook. 2014;1–29.

5. Reynoldson JA, Behnke JM, Pallant LJ, Macnish MG, Gilbert F, Giles S, Spargo RJ, Thompson RC. Failure of pyrantel in treatment of human hookworm infections (*Ancylostoma duodenale*) in the Kimberley region of north west Australia. Acta Trop. 1997;68(3):301–312.

6. De Clercq D, Sacko M, Behnke J, Gilbert F, Dorny P, Vercruysse J. Failure of mebendazole in treatment of human hookworm infections in the southern region of Mali. Am J Trop Med Hyg. 1997;57(1):25–30.

7. Osei-Atweneboana MY, Awadzi K, Attah SK, Boakye DA, Gyapong JO, Prichard RK. Phenotypic evidence of emerging ivermectin resistance in *Onchocerca volvulus*. PLoS Negl Trop Dis. 2011; doi:10.1371/journal.pntd.0000998.

8. Charlier J, Morgan ER, Rinaldi L, van Dijk J, Demeler J, Höglund J, Hertzberg H, Van Ranst B, Hendrickx G, Vercruysse J, Kenyon F. Practices to optimise gastrointestinal nematode control on sheep, goat and cattle farms in Europe using targeted (selective) treatments. Vet Rec. 2014;175(10):250–255.

9. Pulavarty A, Egan A, Karpinska A, Horgan K, Kakouli-Duarte T. Plant parasitic nematodes: a review on their behaviour, host interaction, management approaches and their occurrence in two sites in the Republic of Ireland. Plants (Basel). 2021; doi:10.3390/plants10112352.

10. Coppieters W, Mes TH, Druet T, Farnir F, Tamma N, Schrooten C, Cornelissen AW, Georges M, Ploeger HW. Mapping QTL influencing gastrointestinal nematode burden in Dutch Holstein-Friesian dairy cattle. BMC Genomics. 2009; doi:10.1186/1471-2164-10-96.

11. Scott I, Pomroy WE, Kenyon PR, Smith G, Adlington B, Moss A. Lack of efficacy of monepantel against *Teladorsagia circumcincta* and *Trichostrongylus colubriformis*. Vet Parasitol. 2013;198(1-2):166–171.

12. Burns AR, Luciani GM, Musso G, Bagg R, Yeo M, Zhang Y, Rajendran L, Glavin J, Hunter R, Redman E, Stasiuk S, Schertzberg M, Angus MG, Caffrey CR, Cutler SR, Tyers M, Giaever G, Nislow C, Fraser AG, MacRae CA, Gilleard J, Roy PJ. *Caenorhabditis elegans* is a useful model for anthelmintic discovery. Nat Commun. 2015; doi:10.1038/ncomms8485.

13. Risi G, Aguilera E, Ladós E, Suárez G, Carrera I, Álvarez G, Salinas G. *Caenorhabditis elegans* infrared-based motility assay identified new hits for nematicide drug development. Vet Sci. 2019; doi:10.3390/vetsci6010029.

14. Blanco MG, Vela Gurovic MS, Silbestri GF, Garelli A, Giunti S, Rayes D, De Rosa MJ. Diisopropylphenyl-imidazole (DII): A new compound that exerts anthelmintic activity through novel molecular mechanisms. PLoS Negl Trop Dis. 2018; doi:10.1371/journal.pntd.0007021.

15. Partridge FA, Brown AE, Buckingham SD, Willis NJ, Wynne GM, Forman R, Else KJ, Morrison AA, Matthews JB, Russell AJ, Lomas DA, Sattelle DB. An automated high-throughput system for phenotypic screening of chemical libraries on *C. elegans* and parasitic nematodes. Int J Parasitol Drugs Drug Resist. 2018;8(1):8–21.

16. Medicines for Malaria Venture: MMV Open stimulating research into neglected and pandemic diseases. https://www.mmv.org/newsroom/interviews/mmv-open-stimulating-research-neglected-and-pandemic-diseases (2020). Accessed 1 Dec 2023.

17. Kanatani S, Elahi R, Kanchanabhogin S, Vartak N, Tripathi AK, Prigge ST, Sinnis P. Screening the pathogen box for inhibition of *Plasmodium falciparum* sporozoite motility reveals a critical role for kinases in transmission stages. Antimicrob Agents Chemother. 2022; doi:10.1128/aac.00418-22.

18. Paiardini A, Bamert RS, Kannan-Sivaraman K, Drinkwater N, Mistry SN, Scammells PJ, McGowan S. Screening the medicines for malaria venture "malaria box" against the plasmodium falciparum aminopeptidases, M1, M17 and M18. PLoS One. 2015; doi:10.1371/journal.pone.0115859.

19. Radhakrishnan A, Brown CM, Guy CS, Cooper C, Pacheco-Gomez R, Stansfeld PJ, Fullam E. Interrogation of the Pathogen Box reveals small molecule ligands against the mycobacterial trehalose transporter LpqY-SugABC. RSC Med Chem. 2022;13(10):1225–1233.

20. Jefferson T, McShan D, Warfield J, Ogungbe IV. Screening and identification of inhibitors of *trypanosoma brucei* cathepsin l with antitrypanosomal activity. Chem Biol Drug Des. 2016;87(1):154–158.

21. Boyom FF, Fokou PV, Tchokouaha LR, Spangenberg T, Mfopa AN, Kouipou RM, Mbouna CJ, Donfack VF, Zollo PH. Repurposing the open access malaria box to discover potent inhibitors of *Toxoplasma gondii* and *Entamoeba histolytica*. Antimicrob Agents Chemother. 2014;58(10):5848–5854.

22. Ingram-Sieber K, Cowan N, Panic G, Vargas M, Mansour NR, Bickle QD, Wells TN, Spangenberg T, Keiser J. Orally active antischistosomal early leads identified from the open access malaria box. PLoS Negl Trop Dis. 2014; doi:10.1371/journal.pntd.0002610.

23. Maccesi M, Aguiar PHN, Pasche V, Padilla M, Suzuki BM, Montefusco S, Abagyan R, Keiser J, Mourão MM, Caffrey CR. Multi-center screening of the Pathogen Box collection for schistosomiasis drug discovery. Parasit Vectors. 2019; doi:10.1186/s13071-019-3747-6.

24. Preston S, Jiao Y, Jabbar A, McGee SL, Laleu B, Willis P, Wells TNC, Gasser RB. Screening of the ’Pathogen Box’ identifies an approved pesticide with major anthelmintic activity against the barber’s pole worm. Int J Parasitol Drugs Drug Resist. 2016;6(3):329–334.

25. Stiernagle T. Maintenance of C. elegans. WormBook. 2006;1–11.

26. Simonetta SH, Golombek DA. An automated tracking system for *Caenorhabditis elegans* locomotor behavior and circadian studies application. J Neurosci Methods. 2007;161(2):273–280.

27. Ahmed SA, Gogal RM Jr, Walsh JE. A new rapid and simple non-radioactive assay to monitor and determine the proliferation of lymphocytes: an alternative to [3H]thymidine incorporation assay. J Immunol Methods. 1994;170(2):211–224.

28. Sandhu A, Badal D, Sheokand R, Tyagi S, Singh V. Specific collagens maintain the cuticle permeability barrier in *Caenorhabditis elegans*. Genetics. 2021; doi:10.1093/genetics/iyaa047.

29. Lindblom TH, Dodd AK. Xenobiotic detoxification in the nematode *Caenorhabditis elegans*. Journal of experimental zoology. J Exp Zool A Comp Exp Biol. 2006;305(9):720–730.

30. Geary TG, Moreno Y. Macrocyclic lactone anthelmintics: spectrum of activity and mechanism of action. Curr Pharm Biotechnol. 2012;13(6):866–872.

31. Yu SJ. The toxicology and biochemistry of insecticides. 2nd ed. CRC Press; 2014.

32. Liu L, Caffrey CR. Anthelmintics IV: ‘From Discovery to Resistance’; February 3-7, 2020; Santa Monica, CA.

33. Rollins R, Qader M, Gosnell W, Wang C, Cao S, Cowie R. A validated high-throughput method for assaying rat lungworm (*Angiostrongylus cantonensis*) motility when challenged with potentially anthelmintic natural products from Hawaiian fungi. Parasitology. 2022;149(6):765–773.

34. Liu M, Lu JG, Yang MR, Jiang ZH, Wan X, Luyten W. Bioassay-guided isolation of anthelmintic components from *Semen pharbitidis*, and the mechanism of action of pharbitin. Int J Mol Sci. 2022; doi:10.3390/ijms232415739.

35. Tritten L, Silbereisen A, Keiser J. In vitro and in vivo efficacy of monepantel (AAD 1566) against laboratory models of human intestinal nematode infections. PLoS Negl Trop Dis. 2011; doi:10.1371/journal.pntd.0001457.

36. Widmayer SJ, Crombie TA, Nyaanga JN, Evans KS, Andersen EC. *C. elegans* toxicant responses vary among genetically diverse individuals. Toxicology. 2022; doi:10.1016/j.tox.2022.153292.

37. Raghavendra K, Barik TK, Sharma P, Bhatt RM, Srivastava HC, Sreehari U, Dash AP. Chlorfenapyr: a new insecticide with novel mode of action can control pyrethroid resistant malaria vectors. Malar J. 2011;10:16.

38. Kang C, Kim DH, Kim SC, Kim DS. A patient fatality following the ingestion of a small amount of chlorfenapyr. J Emerg Trauma Shock. 2014;7(3):239–241.

39. Chung MJ, Mao YC, Hsu CT, Chung MC, Wang TJ, Yu TM, Liu PY, Fu PK, Hsieh CM. A fatal case of chlorfenapyr poisoning and the therapeutic implications of serum chlorfenapyr and tralopyril levels. Medicina (Kaunas). 2022;58(11):1630.

40. Chien SC, Chien SC, Su YJ. A fatal case of chlorfenapyr poisoning and a review of the literature. J Int Med Res. 2022; doi:10.1177/03000605221121965.

41. Ghanim M, Lebedev G, Kontsedalov S, Ishaaya I. Flufenerim, a novel insecticide acting on diverse insect pests: biological mode of action and biochemical aspects. J Agric Food Chem. 2011;59(7):2839–2844.

42. Wolstenholme AJ. Ion channels and receptor as targets for the control of parasitic nematodes. Int J Parasitol Drugs Drug Resist. 2011;1(1):2–13.

43. You H, Liu C, Du X, McManus DP. Acetylcholinesterase and Nicotinic Acetylcholine Receptors in Schistosomes and Other Parasitic Helminths. Molecules. 2017;22(9):1550.

44. Casida JE, Durkin KA. Anticholinesterase insecticide retrospective. Chem Biol Interact. 2013;203(1):221–225.

45. Hancock PM, Walsh M, White SJ, Baugh PJ, Catlow DA. Extraction and determination of the MITINS sulcofuron and flucofuron from Environmental River Water. Analyst. 1998;123(8):1669–1674.

46. York E, McNaughton DA, Roseblade A, Cranfield CG, Gale PA, Rawling T. Structure-Activity Relationship and Mechanistic Studies of Bisaryl Urea Anticancer Agents Indicate Mitochondrial Uncoupling by a Fatty Acid-Activated Mechanism. ACS Chem Biol. 2022;17(8):2065–2073.

47. Srivastava IK, Rottenberg H, Vaidya AB. Atovaquone, a broad spectrum antiparasitic drug, collapses mitochondrial membrane potential in a malarial parasite. J Biol Chem. 1997;272(7):3961–3966.

48. Preston S, Korhonen PK, Mouchiroud L, Cornaglia M, McGee SL, Young ND, Davis RA, Crawford S, Nowell C, Ansell BRE, Fisher GM, Andrews KT, Chang BCH, Gijs MAM, Sternberg PW, Auwerx J, Baell J, Hofmann A, Jabbar A, Gasser RB. Deguelin exerts potent nematocidal activity via the mitochondrial respiratory chain. FASEB J. 2017;31(10):4515–4532.

49. Munjal A, Allam AE. Indomethacin. In: StatPearls. StatPearls Publishing. 2022. https://www.ncbi.nlm.nih.gov/books/NBK555936/. Accessed 1 Dec 2023

50. Simmons DL, Botting RM, Hla T. Cyclooxygenase isozymes: the biology of prostaglandin synthesis and inhibition. Pharmacol Rev. 2004;56(3):387–437.

51. Hoang HD, Prasain JK, Dorand D, Miller MA. A heterogeneous mixture of F-series prostaglandins promotes sperm guidance in the *Caenorhabditis elegans* reproductive tract. PLoS Genet. 2013; doi:10.1371/journal.pgen.1003271.

52. Tiwary E, Hu M, Miller MA, Prasain JK. Signature profile of cyclooxygenase-independent F2 series prostaglandins in *C. elegans* and their role in sperm motility. Sci Rep. 2019; doi:10.1038/s41598-019-48062-y.

53. Abdellatif KRA, Abdelall EKA, Elshemy HAH, El-Nahass ES, Abdel-Fattah MM, Abdelgawad YYM. New indomethacin analogs as selective COX-2 inhibitors: Synthesis, COX-1/2 inhibitory activity, anti-inflammatory, ulcerogenicity, histopathological, and docking studies. Arch Pharm. 2021; doi:10.1002/ardp.202000328.

